# Identification of novel, clinically correlated autoantigens in the monogenic autoimmune syndrome APS1 by PhIP-Seq

**DOI:** 10.1101/2020.01.20.913186

**Authors:** Sara E. Vazquez, Elise M. N. Ferré, David W. Scheel, Sara Sunshine, Brenda Miao, Caleigh Mandel-Brehm, Zoe Quandt, Alice Y. Chan, Mickie Cheng, Michael S. German, Michail S. Lionakis, Joseph L. DeRisi, Mark S. Anderson

**Author notes:** These authors contributed equally (Joseph L. DeRisi, Mark S. Anderson). Address Correspondence to: Mark S. Anderson, or Joseph L. DeRisi.

## Abstract

The identification of autoantigens remains a critical challenge for understanding and treating autoimmune diseases. Autoimmune polyendocrine syndrome type 1 (APS1), a rare monogenic form of autoimmunity, presents as widespread autoimmunity with T and B cell responses to multiple organs. Importantly, autoantibody discovery in APS1 can illuminate fundamental disease pathogenesis, and many of the antigens found in APS1 extend to common autoimmune diseases. Here, we performed proteome-wide programmable phage-display (PhIP-Seq) on sera from an APS1 cohort and discovered multiple common antibody targets. These novel autoantigens exhibit tissue-restricted expression, including expression in enteroendocrine cells and dental enamel. Using detailed clinical phenotyping, we find novel associations between autoantibodies and organ-restricted autoimmunity, including between anti-KHDC3L autoantibodies and premature ovarian insufficiency, and between anti-RFX6 autoantibodies and diarrheal-type intestinal dysfunction. Our study highlights the utility of PhIP-Seq for interrogating antigenic repertoires in human autoimmunity and the importance of antigen discovery for improved understanding of disease mechanisms.

## INTRODUCTION

Autoimmune Polyglandular syndrome type 1 (APS1) or Autoimmune Polyglandular-Candidiasis-Ectodermal Dystrophy (APECED; OMIM #240300) is an autoimmune syndrome caused by monogenic mutations in the *AIRE* gene that result in defects in AIRE-dependent T cell education in the thymus (Aaltonen et al., 1997; Anderson, 2002; Conteduca et al., 2018; Malchow et al., 2016; Nagamine et al., 1997). As a result, people with APS1 develop autoimmunity to multiple organs, including endocrine organs, skin, gut, and lung (Ahonen et al., 1990; Ferré et al., 2016; Söderbergh et al., 2004). Although the majority of APS1 autoimmune manifestations are thought to be primarily driven by autoreactive T cells, people with APS1 also possess autoreactive B cells and corresponding high-affinity autoantibody responses (DeVoss et al., 2008; Gavanescu et al., 2008; Meyer et al., 2016; Sng et al., 2019). These autoantibodies likely derive from germinal center reactions driven by self-reactive T cells, resulting in mirroring of autoantigen identities between the T and B cell compartments (Lanzavecchia, 1985; Meyer et al., 2016).

Identification of the specificity of autoantibodies in autoimmune diseases is important for understanding underlying disease pathogenesis and for identifying those at risk for disease (Rosen & Casciola-Rosen, 2014). However, despite the long-known association of autoantibodies with specific diseases in both monogenic and sporadic autoimmunity, many autoantibody specificities remain undiscovered. Challenges in antigen identification include the weak affinity of some autoantibodies for their target antigen, as well as rare or low expression of the target antigen. One approach to overcome some of these challenges is to interrogate autoimmune patient samples with particularly high affinity autoantibodies. Indeed, such an approach identified GAD65 as a major autoantigen in type 1 diabetes by using sera from people with Stiff Person Syndrome (OMIM #184850), who harbor high affinity autoantibodies (Baekkeskov et al., 1990). We reasoned that PhIP-Seq interrogation of APS1, a defined monogenic autoimmune syndrome with a broad spectrum of high affinity autoantibodies, would likely yield clinically meaningful targets – consistent with previously described APS1 autoantibody specificities that exhibit strong, clinically useful associations with their respective organ-specific diseases (Alimohammadi et al., 2008, 2009; Ferré et al., 2019; Landegren et al., 2015; Popler et al., 2012; Puel et al., 2010; Shum et al., 2013; Söderbergh et al., 2004; Winqvist et al., 1993).

The identification of key B cell autoantigens in APS1 has occurred most commonly through candidate-based approaches and by whole-protein microarrays. For example, lung antigen BPIFB1 autoantibodies, which are used to assess people with APS1 for risk of interstitial lung disease, were discovered first in *Aire*-deficient mice using a combination of targeted immunoblotting, tissue microscopy, and mass spectrometry (Shum et al., 2013, 2009). Recently, there have been rapid advances in large platform approaches for antibody screening; these platforms can overcome problems of antigen abundance by simultaneously screening the majority of proteins from the human genome in an unbiased fashion (Jeong et al., 2012; Larman et al., 2011; Sharon & Snyder, 2014; Zhu et al., 2001). In particular, a higher-throughput antibody target profiling approach utilizing a fixed protein microarray technology (ProtoArray) has enabled detection of a wider range of proteins targeted by autoantibodies directly from human serum (Fishman et al., 2017; Landegren et al., 2016; Meyer et al., 2016). Despite initial success of this technology in uncovering shared antigens across APS1 cohorts, it is likely that many shared antigens remain to be discovered, given that these arrays do not encompass the full coding potential of the proteome.

Here, we took an alternate approach to APS1 antigen discovery by employing Phage Immunoprecipitation-Sequencing (PhIP-Seq) based on an established proteome-wide tiled library (Larman et al., 2011; O’Donovan et al., 2018). This approach possesses many potential advantages over previous candidate-based and whole-protein fixed array approaches, including (1) expanded, proteome-wide coverage (including alternative splice forms) with 49 amino acid (AA) peptide length and 24AA resolution tiling, (2) reduced volume requirement for human serum, and (3) high-throughput, sequencing based output (Larman et al., 2011; O’Donovan et al., 2018). Of note, the PhIP-Seq investigation of autoimmune diseases of the central nervous system, including paraneoplastic disease, has yielded novel and specific biomarkers of disease (Larman et al., 2011; Mandel-Brehm et al., 2019; O’Donovan et al., 2018).

Using a PhIP-Seq autoantibody survey, we identify a collection of novel APS1 autoantigens as well as numerous known, literature-reported APS1 autoantigens. We orthogonally validate seven novel autoantigens including RFX6, KHDC3L, and ACP4, all of which exhibit tissue-restricted expression (Jeong et al., 2012; Larman et al., 2011; Sharon & Snyder, 2014; H. Zhu et al., 2001). Importantly, these novel autoantigens may carry important implications for poorly understood clinical manifestations such as intestinal dysfunction, ovarian insufficiency, and tooth enamel hypoplasia, where underlying cell-type specific antigens have remained elusive. Together, our results demonstrate the applicability of PhIP-Seq to antigen discovery, substantially expand the spectrum of known antibody targets and clinical associations in APS1, and point towards novel specificities that can be targeted in autoimmunity.

## RESULTS

### Investigation of APS1 serum autoantibodies by PhIP-Seq

Individuals with APS1 develop autoantibodies to many known protein targets, some of which exhibit tissue-restricted expression and have been shown to correlate with specific autoimmune disease manifestations. However, the target proteins for many of the APS1 tissue-specific manifestations remain enigmatic. To this end, we employed a high-throughput, proteome-wide programmable phage display approach (PhIP-Seq) to query the antibody target identities within serum of people with APS1 (Larman et al., 2011; O’Donovan et al., 2018). The PhIP-Seq technique leverages large scale oligo production and efficient phage packaging and expression to present a tiled-peptide representation of the proteome displayed on T7 phage. Here, we utilize a phage library that we previously designed and deployed for investigating paraneoplastic autoimmune encephalitis (Mandel-Brehm et al., 2019; O’Donovan et al., 2018). The library itself contains approximately 700,000 unique phage, each displaying a 49 amino acid proteome segment. As previously described, phage were immunoprecipitated using human antibodies bound to protein A/G beads. In order to increase sensitivity and specificity for target proteins, eluted phage were used for a further round of amplification and immunoprecipitation. DNA was then extracted from the final phage elution, amplified and barcoded, and subjected to Next-Generation Sequencing (**Figure 1A**). Finally, sequencing counts were normalized across samples to correct for variability in sequencing depth, and the fold-change of each gene was calculated (comprised of multiple unique tiling phage) as compared to mock IPs in the absence of human serum (further details of the protocol can be found in the methods section).

**Figure 1.**
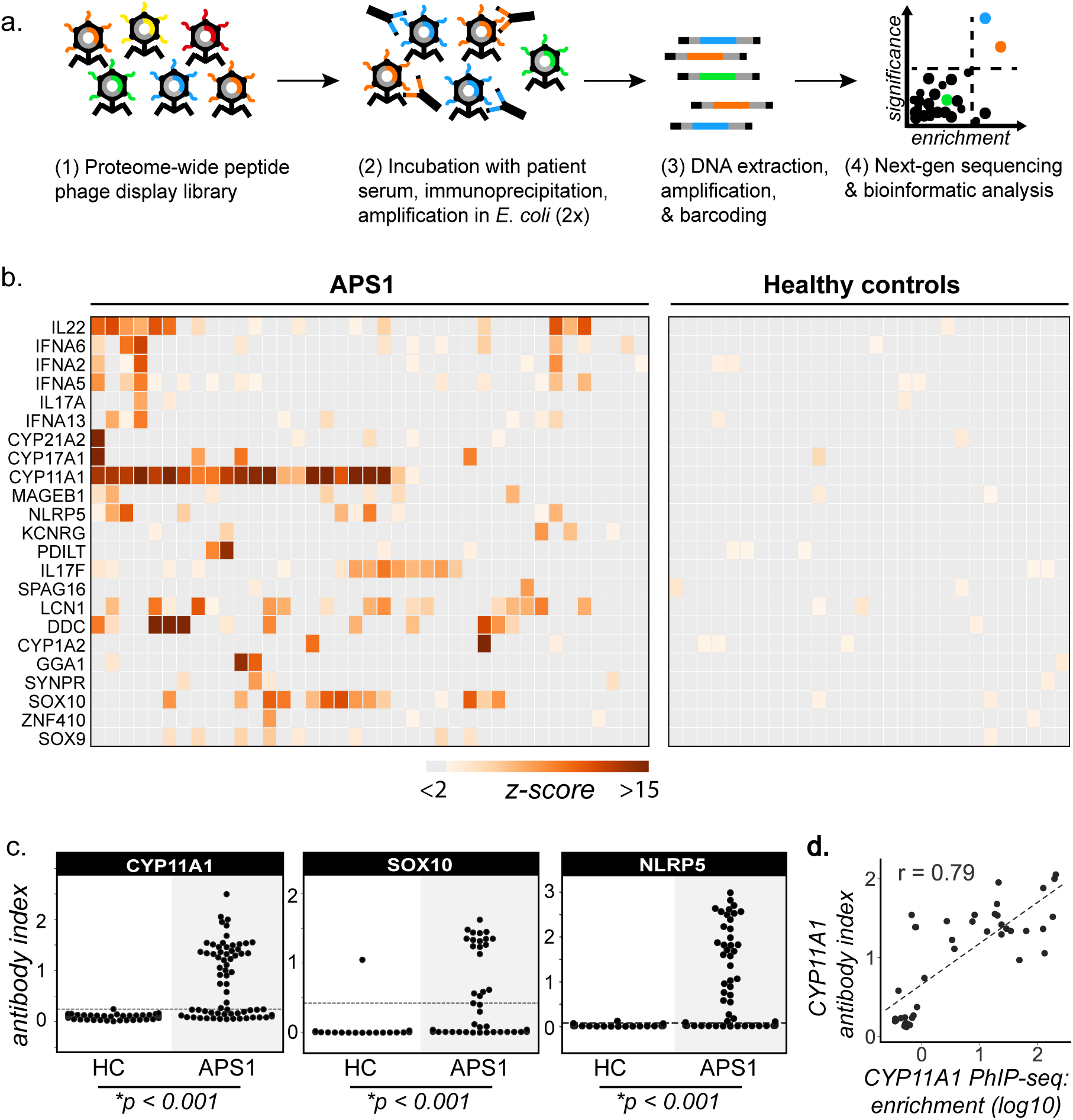
PhIP-Seq identifies literature-reported autoantigens in APS1. **A.** Overview of PhIP-Seq experimental workflow. **B.** PhIP-Seq identifies known autoantibody targets in APS1. Hierarchically clustered (Pearson) z-scored heatmap of literature reported autoantigens with 10-fold or greater signal over mock-IP in at least 2/39 APS1 sera and in 0/28 non-APS1 control sera. See also Figure 1: Figure Supplement 1. **C**. Radioligand binding assay (RLBA) orthogonal validation of literature-reported antigens CYP11A1, SOX10, and NLRP5 within the expanded cohort of APS1 (n = 67) and non-APS1 controls (n = 61); p-value was calculated across all samples using a Mann-Whitney U test. **D**. CYP11A1 RLBA antibody index and CYP11A1 PhIP-Seq enrichment are well correlated (r = 0.79); see also Figure 1: Figure Supplement 2.

From a cohort of 67 APS1 serum samples, a total of 39 samples were subjected to PhIP-Seq investigation, while the remaining 28 samples were obtained at a later time point and reserved for downstream validation experiments (for clinical data, refer to **Supplemental Table 1**). In addition, 28 non-APS1 anonymous blood donor serum samples were subjected to PhIP-Seq, and an additional group of 61 non-APS1 plasma samples were used for downstream validation experiments.

### Detection of literature-reported APS1 autoantigens

PhIP-Seq results were first cross-referenced with previously reported APS1 autoantibody targets (Alimohammadi et al., 2008, 2009; Clemente et al., 1997; Fishman et al., 2017; Hedstrand et al., 2001; Husebye et al., 1997; Kluger et al., 2015; Kuroda et al., 2005; Landegren et al., 2016, 2015; Leonard et al., 2017; Meager et al., 2006; Meyer et al., 2016; Oftedal et al., 2015; Pöntynen et al., 2006; Sansom et al., 2014; Shum et al., 2013, 2009; Söderbergh et al., 2004). To avoid false positives, a conservative set of criteria were used as follows. We required a minimum of 2/39 APS1 samples and 0/28 non-APS1 control samples to exhibit normalized gene counts in the immunoprecipitation (IP) with greater than 10-fold enrichment as compared to the control set of 18 mock-IP (beads, no serum) samples. This simple, yet stringent criteria enabled detection of a total of 23 known autoantibody specificities (**Figure 1B**). Importantly, many of the well-validated APS1 antigens, including specific members of the cytochrome P450 family (CYP1A2, CYP21A1, CYP11A1, CYP17A1), lung disease-associated antigen KCRNG, as well as IL17A, IL17F, and IL22, among others were well represented (**Figure 1B**). In contrast, the diabetes-associated antigens GAD65 and INS did not meet these stringent detection criteria and only weak signal was detected to many of the known interferon autoantibody targets known to be present in many people with APS1, perhaps due to the conformational nature of these autoantigens (**Figure 1B & Figure 1: Figure Supplement 1**) (Björk et al., 1994; Meager et al., 2006; Meyer et al., 2016; Wolff et al., 2013; Ziegler et al., 1996).

Three known autoantigens that were prevalent within our cohort were selected to determine how PhIP-Seq performed against an orthogonal whole protein-based antibody detection assay. A radioligand binding assay (RLBA) was performed by immunoprecipitating *in vitro* transcribed and translated S35-labeled proteins CYP11A1, SOX10, and NLRP5 with APS1 serum (Alimohammadi et al., 2008; Berson et al., 1956; Hedstrand et al., 2001; Winqvist et al., 1993). Importantly, and in contrast to PhIP-Seq, this assay tests for antibody binding to full-length protein (**Figure 1C**). By RLBA, these three antigens were present in and specific to both the initial discovery APS1 cohort (n=39) as well as the expanded validation cohort (n = 28), but not the non-APS1 control cohort (n = 61). Together, these results demonstrate that PhIP-Seq detects known APS1 autoantigens and that PhIP-Seq results validate well in orthogonal whole protein-based assays.

To determine whether the PhIP-Seq APS1 dataset could yield higher resolution information on antigenic peptide sequences with respect to previously reported targets, the normalized enrichments of all peptides belonging to known disease-associated antigens CYP11A1 and SOX10 were mapped across the full length of their respective proteins (**Figure 1: Figure Supplement 2**). The antigenic regions within these proteins were observed to be similar across all samples positive for anti-CYP11A1 and anti-SOX10 antibodies, respectively (**Figure 1: Figure Supplement 2**) suggesting peptide-level commonalities and convergence among the autoreactive antibody repertoires across individuals. These data suggest that people with APS1 often target similar, but not identical protein regions.

### Identification of novel APS1 autoantigens

Having confirmed that PhIP-Seq analysis of APS1 sera detected known antigens, the same data were then investigated for the presence of novel, previously uncharacterized APS1 autoantigens. We applied the same positive hit criteria as described for known antigens, and additionally increased the required number of positive APS1 samples to 3/39 to impose a stricter limit on the number of novel candidate autoantigens. This yielded a list of 82 genes, which included 10 known antigens and 72 putative novel antigens (**Figure 2**).

**Figure 2.**
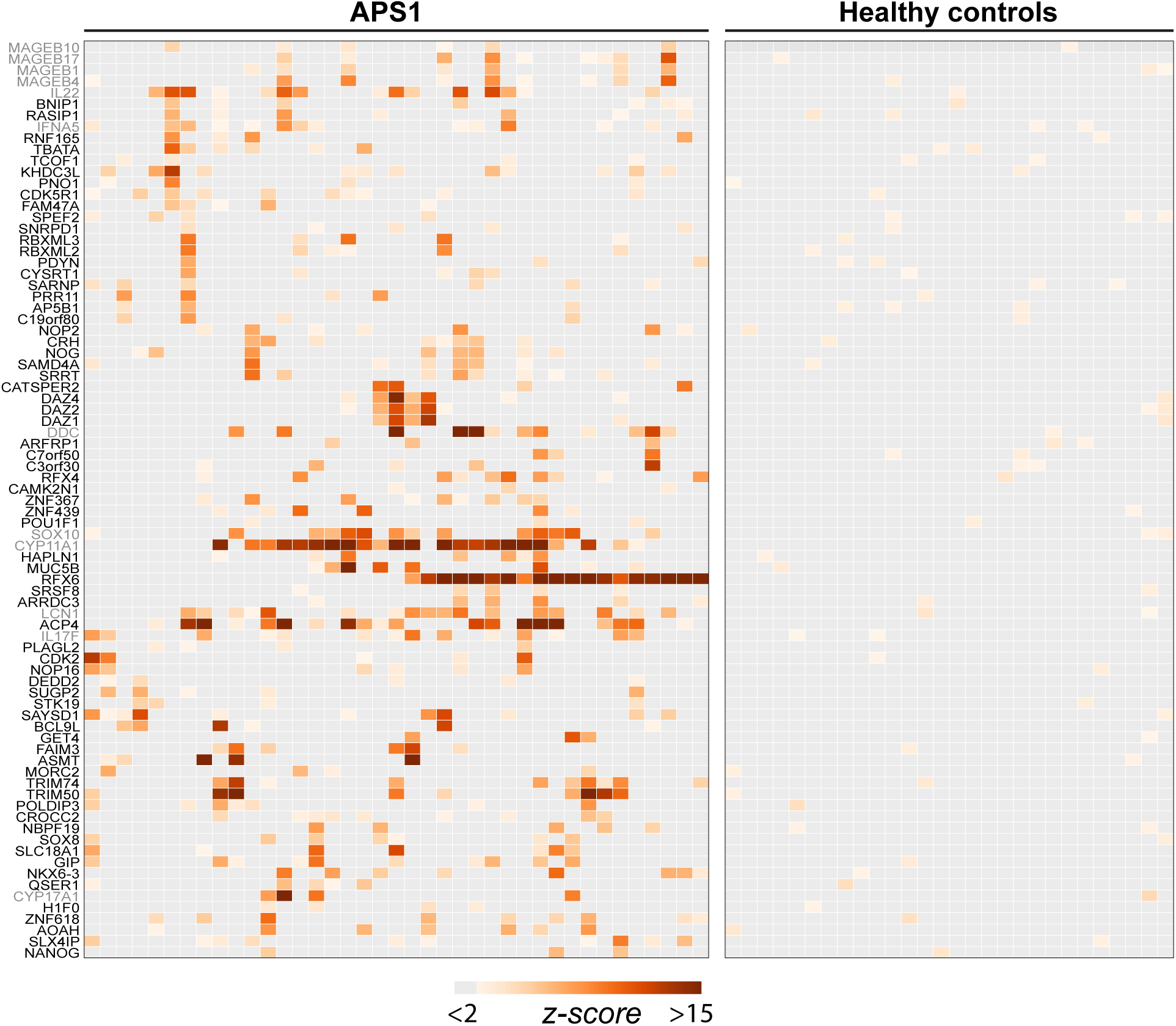
PhIP-Seq identifies novel (and known) antigens across multiple APS1 sera. **A**. Hierarchically clustered (Pearson) Z-scored heatmap of all genes with 10-fold or greater signal over mock-IP in at least 3/39 APS1 sera and in 0/28 non-APS1 sera. Black labeled antigens (n=69) are potentially novel and grey labeled antigens (n=12) are previously literature-reported antigens. See also Figure 2: Figure Supplement 1.

The most commonly held hypotheses regarding the nature and identity of proteins targeted by the aberrant immune response in APS1 are that targeted proteins (1) tend to exhibit AIRE-dependent thymic expression and (2) have restricted expression to one or few peripheral organs and tend not to be widely or ubiquitously expressed. We investigated whether our novel antigens were also preferentially tissue-restricted. In order to systematically address this question, tissue-specific RNA expression was assessed using a consensus expression dataset across 74 cell types and tissues (Uhlen et al., 2015). For each gene, the ratio of expression in the highest tissue as compared to the sum of expression across all tissues was calculated, resulting in higher ratios for those mRNAs with greater degrees of tissue-restriction. Using this approach, the mean tissue-specificity ratio of the 82 PhIP-Seq positive antigens was increased by approximately 1.5-fold (p=0.0017) as compared to the means from iterative sampling of 82 genes (**Figure 2: Figure Supplement 1**).

### Identification of novel antigens common to many individuals

Identified autoantigens were ranked by frequency within the cohort. Five antigens were positive in ten or more APS1 samples, including two novel antigens. In addition, the majority of antigens found in 4 or more APS1 sera were novel (**Figure 3A**). Five of the most frequent novel antigens were selected for subsequent validation and follow-up. These included RFX6, a transcription factor implicated in pancreatic and intestinal pathology (Patel et al., 2017; S. B. Smith et al., 2010); ACP4, an enzyme implicated in dental enamel hypoplasia (Choi et al., 2016; Seymen et al., 2016; C. E. Smith et al., 2017); KHDC3L, a protein with oocyte-restricted expression (Li et al., 2008; Zhang et al., 2018; K. Zhu et al., 2015); NKX6-3, a gastrointestinal transcription factor (Alanentalo et al., 2006); and GIP, a gastrointestinal peptide involved in intestinal motility and energy homeostasis (Adriaenssens et al., 2019; Moody et al., 1984; Pederson & McIntosh, 2016). Several less frequent (but still shared) novel antigens were also chosen for validation, including ASMT, a pineal gland enzyme involved in melatonin synthesis (Ackermann et al., 2006; Rath et al., 2016); and PDX1, an intestinal and pancreatic transcription factor (Holland et al., 2002; Stoffers et al., 1997) (**Figure 3A**). Of note, this group of seven novel antigens all exhibited either tissue enriched, tissue enhanced, or group enhanced expression according to the Human Protein Atlas database (Uhlen et al., 2015) (**Supplemental Table 2**). Using a whole-protein radiolabeled binding assay (RLBA) for validation, all seven proteins were immunoprecipitated by antibodies in both the PhIP-Seq APS1 discovery cohort (n=39), as well as in the validation cohort of APS1 sera that had not been interrogated by PhIP-Seq (n=28). Whereas an expanded set of non-APS1 controls (n=61) produced little to no immunoprecipitation signal by RLBA as compared to positive control antibodies (low antibody index), APS1 samples yielded significant immunoprecipitation signal enrichment for each whole protein assay (high antibody index) (**Figure 3B & Supplemental Table 3**).

**Figure 3.**
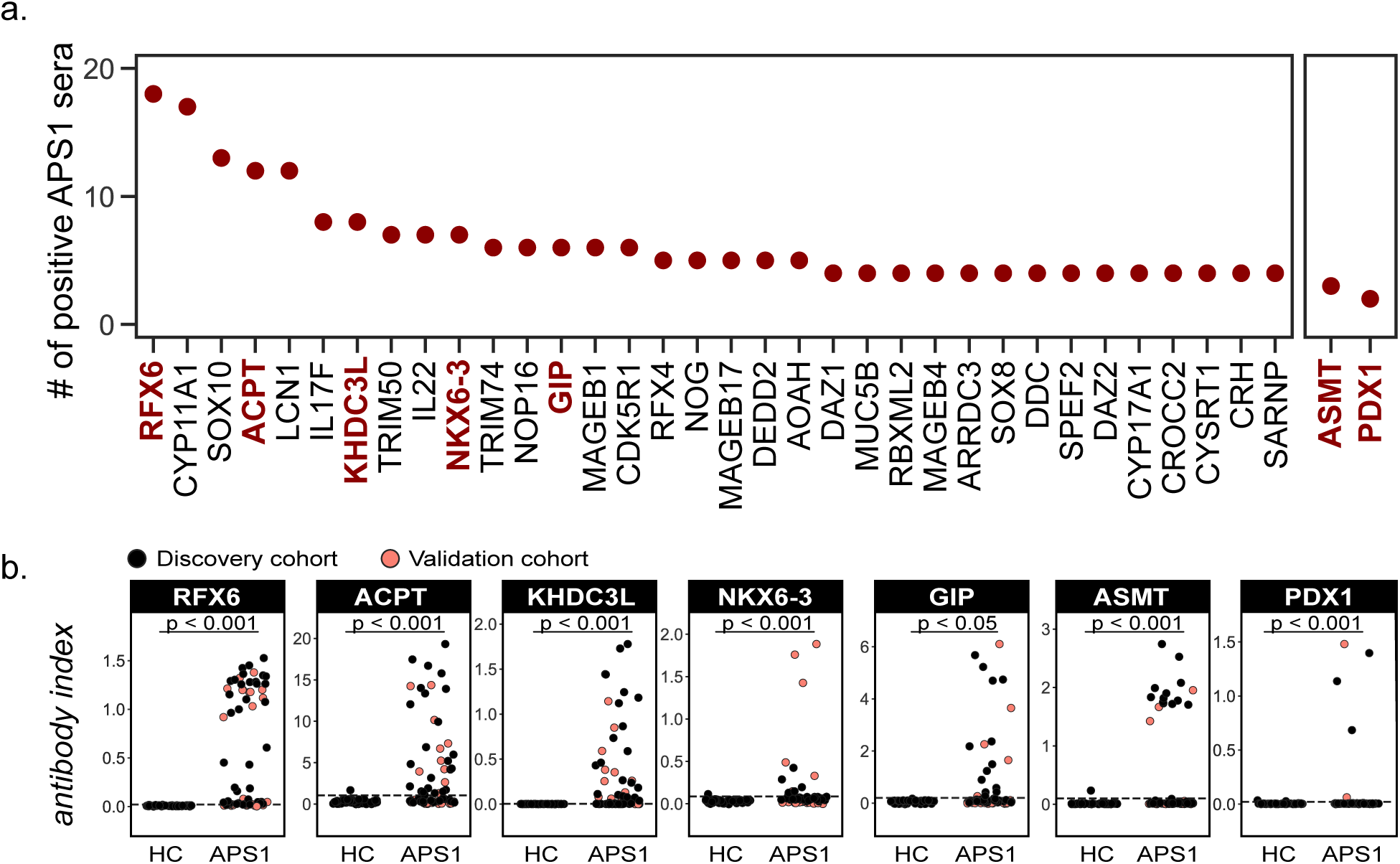
Novel PhIP-Seq autoantigens are shared across multiple APS1 samples and validate in whole protein binding assays. **A.** Graph of the PhIP-seq autoantigens from Figure 2 that were shared across the highest number of individual APS1 sera (left panel). ASMT and PDX1 were positive hits in 3 and 2 sera, respectively, but are known to be highly tissue specific (right panel). Genes in red were chosen for validation in whole protein binding assay. **B.** Validation of novel PhIP-Seq antigens by radiolabeled binding assay, with discovery cohort (black, n_APS1_ = 39), validation cohort (light red, n_APS1_ = 28) and non-APS1 control cohort (n_HC_ = 61). P-value was calculated across all samples using a Mann-Whitney U test. See also Figure 3: Figure Supplement 1.

The comparison of PhIP-Seq data to the results from the RLBAs (n=39, discovery cohort only) yielded positive correlations between the two datasets (r = 0.62-0.95; **Figure 3: Figure Supplement 1**). Notably, for some antigens, such as NLRP5, and particularly for ASMT, the RLBA results revealed additional autoantibody-positive samples not detected by PhIP-Seq (**Figure 3B & Figure 3: Figure Supplement 1 & Figure 1: Figure Supplement 2**).

### Autoantibody-disease associations for both known and novel antigens

Because the individuals in this APS1 cohort have been extensively phenotyped for 24 clinical manifestations, the PhIP-Seq APS1 data was queried for phenotypic associations. Several autoantibody specificities, both known and novel, were found to possess highly significant associations with several clinical phenotypes (**Figure 4 & Figure 4: Figure Supplement 1**). Among these were the associations of KHDC3L with ovarian insufficiency, RFX6 with diarrheal-type intestinal dysfunction, CYP11A1 (also known as cholesterol side chain cleavage enzyme) with adrenal insufficiency (AI), and SOX10 with vitiligo (**Figure 4**). Strikingly, anti-CYP11A1 antibodies are present in AI and are known to predict disease development (Betterle et al., 2002; Obermayer-Straub et al., 2000; Winqvist et al., 1993). Similarly, antibodies to SOX10, a transcription factor involved in melanocyte differentiation and maintenance, have been previously shown to correlate with the presence of autoimmune vitiligo (Hedstrand et al., 2001).

**Figure 4.**
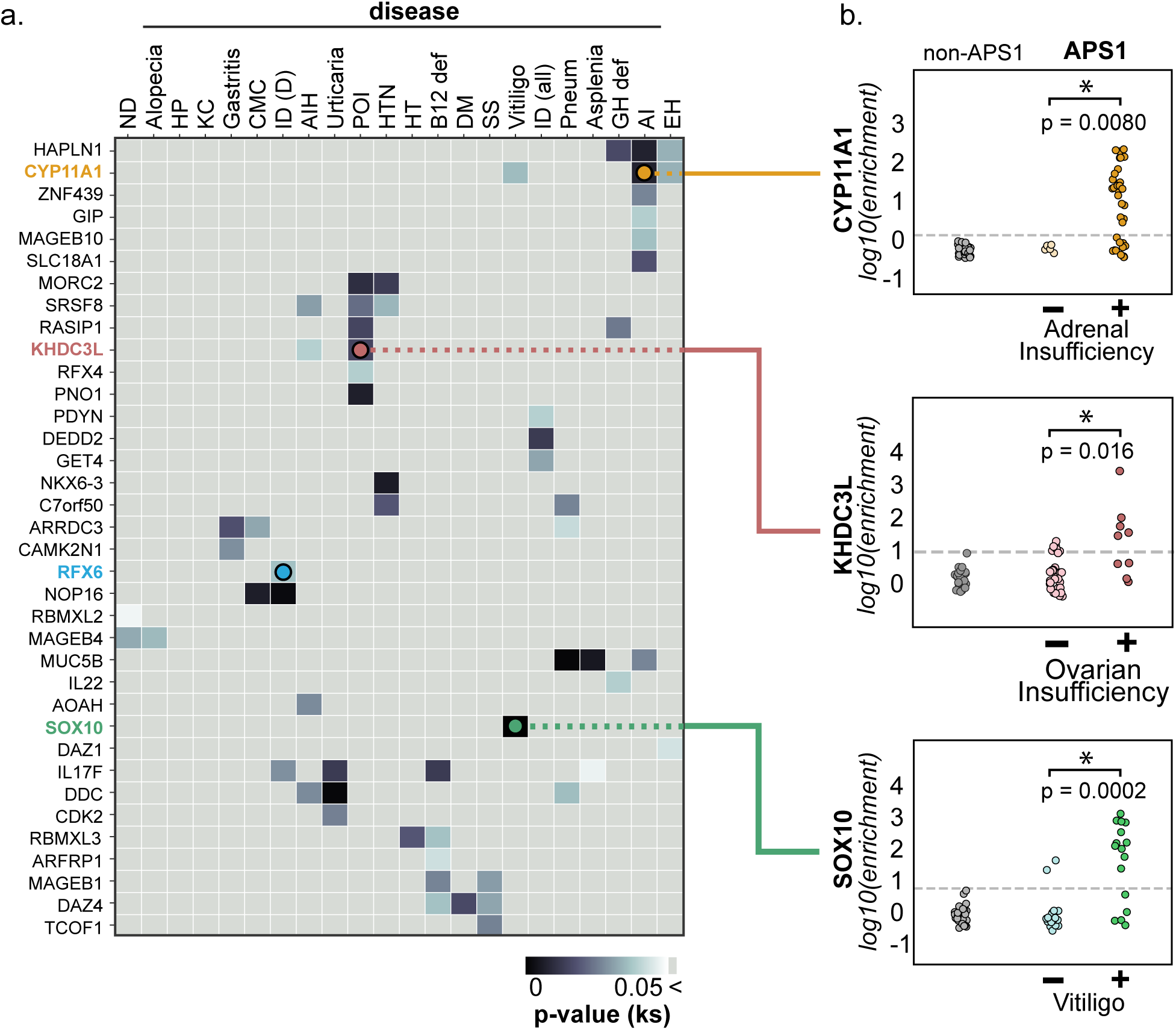
PhIP-Seq reproduces known clinical associations with anti-CYP11A1 and anti-SOX10 antibodies. **A.** Heatmap of p-values (Kolmogorov-Smirnov testing) for differences in gene enrichments for individuals with versus without each clinical phenotype. Significant p-values in the negative direction (where mean PhIP-Seq enrichment is higher in individuals without disease) are masked (colored >0.05). See also Figure 4: Figure Supplement 1. **B.** Anti-CYP11A1 PhIP-Seq enrichments are significantly different between APS1 patients with and without adrenal insufficiency (top panel; Kolmogorov-Smirnov test). Anti-SOX10 PhIP-Seq enrichments are significantly different between APS1 patients with and without Vitiligo (bottom panel). Anti-KHDC3L PhIP-Seq enrichments are significantly different between APS1 patients with and without ovarian insufficiency (middle panel). See also Figure 4: Figure Supplement 2. ND, nail dystrophy. HP, hypoparathyroidism. KC, keratoconjunctivitis. CMC, chronic mucocutaneous candidiasis. ID (D), Intestinal dysfunction (diarrheal-type). AIH, autoimmune hepatitis. POI, primary ovarian insufficiency. HTN, hypertension. HT, hypothyroidism. B12 def, B12 (vitamin) deficiency. DM, diabetes mellitus. SS, Sjogren’s-like syndrome. Pneum, Pneumonitis. GH def, Growth hormone deficiency. AI, Adrenal Insufficiency. EH, (dental) enamel hypoplasia.

### Anti-KHDC3L antibodies in APS1-associated ovarian insufficiency

Primary ovarian insufficiency is a highly penetrant phenotype, with an estimated 60% of females with APS1 progressing to an early, menopause-like state (Ahonen et al., 1990; Ferré et al., 2016). Interestingly, a set of 5 proteins (KHDC3L, SRSF8, PNO1, RASIP1, and MORC2) exhibited a significant association with ovarian insufficiency in this cohort (**Figure 4**). A publicly available RNA-sequencing dataset from human oocytes and supporting granulosa cells of the ovary confirmed that of these 5 genes, only *KHDC3L* exhibited expression levels in female oocytes comparable to the expression levels seen for the known oocyte markers *NLRP5* and *DDX4* (Zhang et al., 2018) (**Figure 4: Figure Supplement 2**). We therefore chose to further investigate the relationship between anti-KHDC3L antibodies and ovarian insufficiency in our cohort (**Figure 5**).

**Figure 5.**
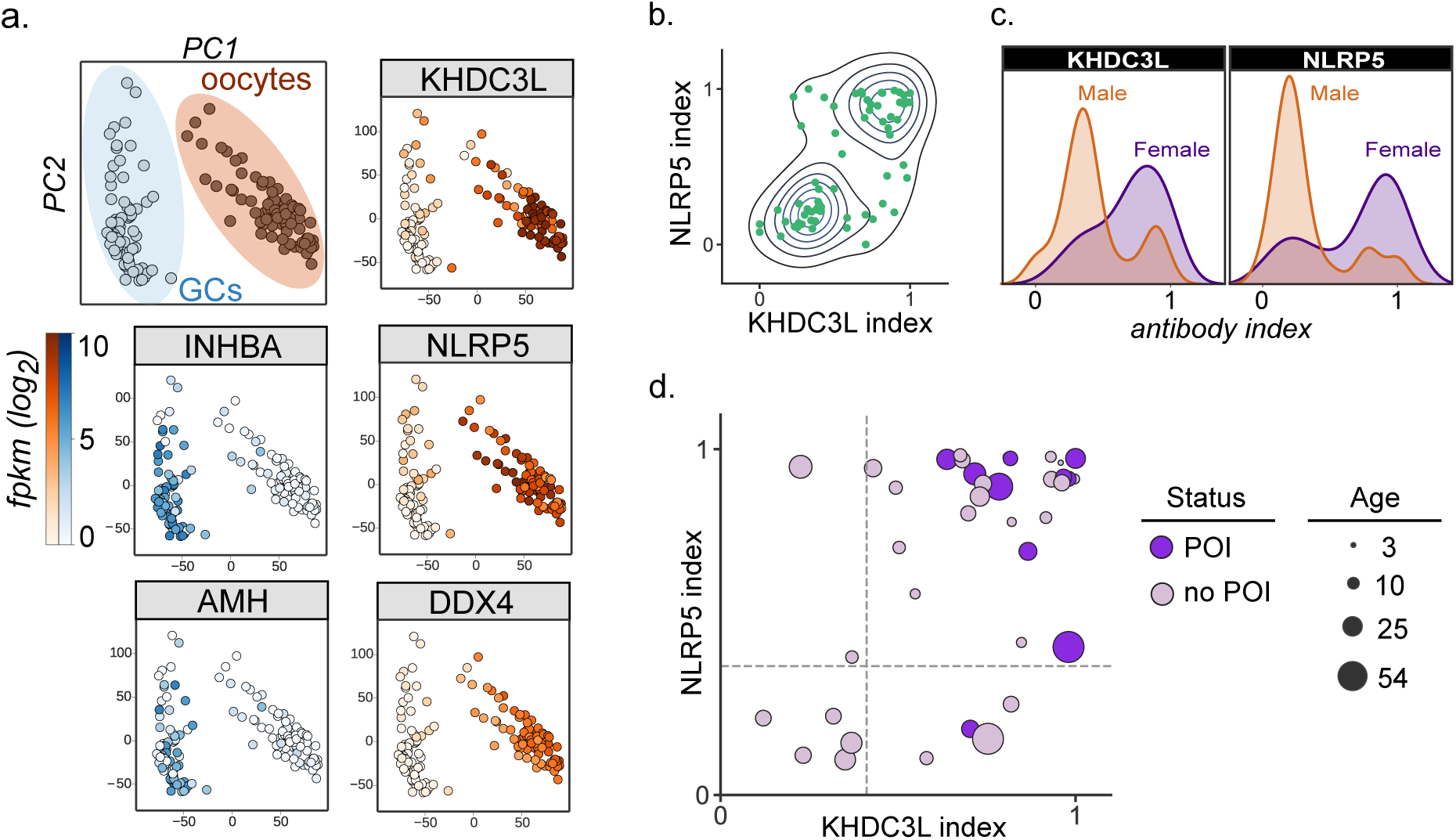
Autoantibodies to oocyte-expressed protein KHDC3L are associated with ovarian insufficiency. **A.** Principle component analysis of transcriptome of single human oocytes (red) and granulosa cells (GCs, blue); data from Zhang et al., Mol Cell 2018. KHDC3L is highly expressed in oocytes, along with binding partner NLRP5 and known oocyte marker DDX4. For comparison, known GC markers INHBA and AMH are primarily expressed in the GC population. **B.** APS1 sera that are positive for one of anti-KHDC3L and anti-NLRP5 autoantibodies tend to also be positive for the other. **C.** Antibody indices for both KHDC3L and NLRP5 are increased in females with APS1. **D.** Antibody indices for females with APS1 by age; All 10 patients with primary ovarian insufficiency (POI) are positive for anti-KHDC3L antibodies. Of note, many of the individuals with anti-KHDC3L antibodies but without POI are younger and therefore cannot be fully evaluated for ovarian insufficiency.

KHDC3L is a well-studied molecular binding partner of NLRP5 within the ovary (Li et al.,2008; K. Zhu et al., 2015). Together, NLRP5 and KHDC3L form part of a critical oocyte-specific molecular complex, termed the subcortical maternal complex (SCMC) (Bebbere et al., 2016; Brozzetti et al., 2015; Li et al., 2008; Liu et al., 2016; K. Zhu et al., 2015). Furthermore, knockout of the *NLRP5* and *KHDC3L* in female mice results in fertility defects, and human genetic mutations in these genes of the SCMC have been linked to infertility and molar pregnancies (Li et al., 2008; Y. Zhang et al., 2018; K. Zhu et al., 2015). Interestingly, previous work established NLRP5 as a parathyroid-specific antigen in APS1, with potential for additional correlation with ovarian insufficiency (Alimohammadi et al., 2008). However, anti-NLRP5 antibodies lack sensitivity for ovarian insufficiency. Importantly, unlike NLRP5, KHDC3L is expressed primarily in the ovary, and thus potentially represents a more oocyte-specific autoantigen (Liu et al., 2016; Virant-Klun et al., 2016; Y. Zhang et al., 2018). Using the dataset from Zhang et. al, we confirmed that KHDC3L, as well as NLRP5 and the known oocyte marker DDX4, are highly expressed within the oocyte population, but not in the supporting granulosa cell types (Y. Zhang et al., 2018) (**Figure 5A**). Interestingly, the majority (64%) of APS1 sera had a concordant status for antibodies to KHDC3L and NLRP5 (**Figure 5B**). Although previous reports did not find a strong gender prevalence within samples positive for anti-NLRP5 antibodies, the mean anti-NLRP5 and anti-KHDC3L antibody signals were increased in females in this cohort (**by 1.6- and 2.1-fold, respectively; Figure 5C**). Finally, all 10 females in the expanded APS1 cohort with diagnosed ovarian insufficiency were also positive for anti-KHDC3L antibodies (**Figure 5D**).

### High prevalence of anti-ACP4 antibodies

Similar to known antigens CYP11A1, SOX10, and LCN1, the novel antigen ACP4 was found to occur at high frequencies in this cohort (**Figure 3A**). ACP4 (acid phosphatase 4) is highly expressed in dental enamel, and familial mutations in the *ACP4* gene result in dental enamel hypoplasia similar to the enamel hypoplasia seen in ∼90% of this APS1 cohort (Seymen et al., 2016; C. E. Smith et al., 2017). Strikingly, 50% of samples were positive for anti-ACP4 antibodies by RLBA, with excellent correlation between RLBA and PhIP-Seq data (**Figure 3B & Figure 3: Figure Supplement 1A**). Consistently, samples from individuals with enamel hypoplasia exhibited a trend towards higher anti-ACP4 antibody signal by RLBA (**Figure 3: Figure Supplement 1B, p = 0.064**).

### High prevalence of anti-RFX6 antibodies

In this cohort, 82% (55/67) of APS1 sera exhibited an RFX6 signal that was at least 3 standard deviations above the mean of non-APS1 control signal due to the extremely low RFX6 signal across all non-APS1 controls by RLBA (**Figure 3B**). Using a more stringent cutoff for RFX6 positivity by RLBA at 6 standard deviations above the mean, 65% of APS1 samples were positive for anti-RFX6 antibodies. RFX6 is expressed in both intestine and pancreas, and loss of function RFX6 variants in humans lead to both intestinal and pancreatic pathology (Gehart et al., 2019; Patel et al., 2017; Piccand et al., 2019; S. B. Smith et al., 2010). Interestingly, across all samples with anti-RFX6 antibodies, the response targeted multiple sites within the protein, suggesting a polyclonal antibody response (**Figure 6: Figure Supplement 1A**).

### Anti-enteroendocrine and anti-RFX6 response in APS1

The extent and frequency of intestinal dysfunction in people with APS1 has only recently been clinically uncovered and reported, and therefore still lacks unifying diagnostic markers as well as specific intestinal target antigen identities (Ferré et al., 2016). This investigation of APS1 sera revealed several antigens that are expressed in the intestine, including RFX6, GIP, PDX1, and NKX6-3. We chose to further study whether autoimmune response to RFX6+ cells in the intestine was involved in APS1-associated intestinal dysfunction. Using a publicly available murine single-cell RNA sequencing dataset of 16 different organs and over 120 different cell types, *RFX6* expression was confirmed to be present in and restricted to pancreatic islets and intestinal enteroendocrine cells (Schaum et al., 2018) (**Figure 6: Figure Supplement 1B & 1C**). Serum from an individual with APS1-associated intestinal dysfunction and anti-RFX6 antibodies was next tested for reactivity against human intestinal enteroendocrine cells, revealing strong nuclear staining that colocalized with ChromograninA (ChgA), a well-characterized marker of intestinal enteroendocrine cells (Goldspink et al., 2018; O’Connor et al., 1983) (**Figure 6A, right panel and inset**). In contrast, enteroendocrine cell staining was not observed from APS1 samples that lacked anti-RFX6 antibodies or from non-APS1 control samples. (**Figure 6A, center & left panels**). Furthermore, serum from samples with anti-RFX6 antibodies stained transfected tissue culture cells expressing RFX6 (**Figure 6B, Figure 6: Figure Supplement 2**). These data support the notion that there exists a specific antibody signature, typified by anti-RFX6 antibodies, associated with enteroendocrine cells in APS1.

**Figure 6.**
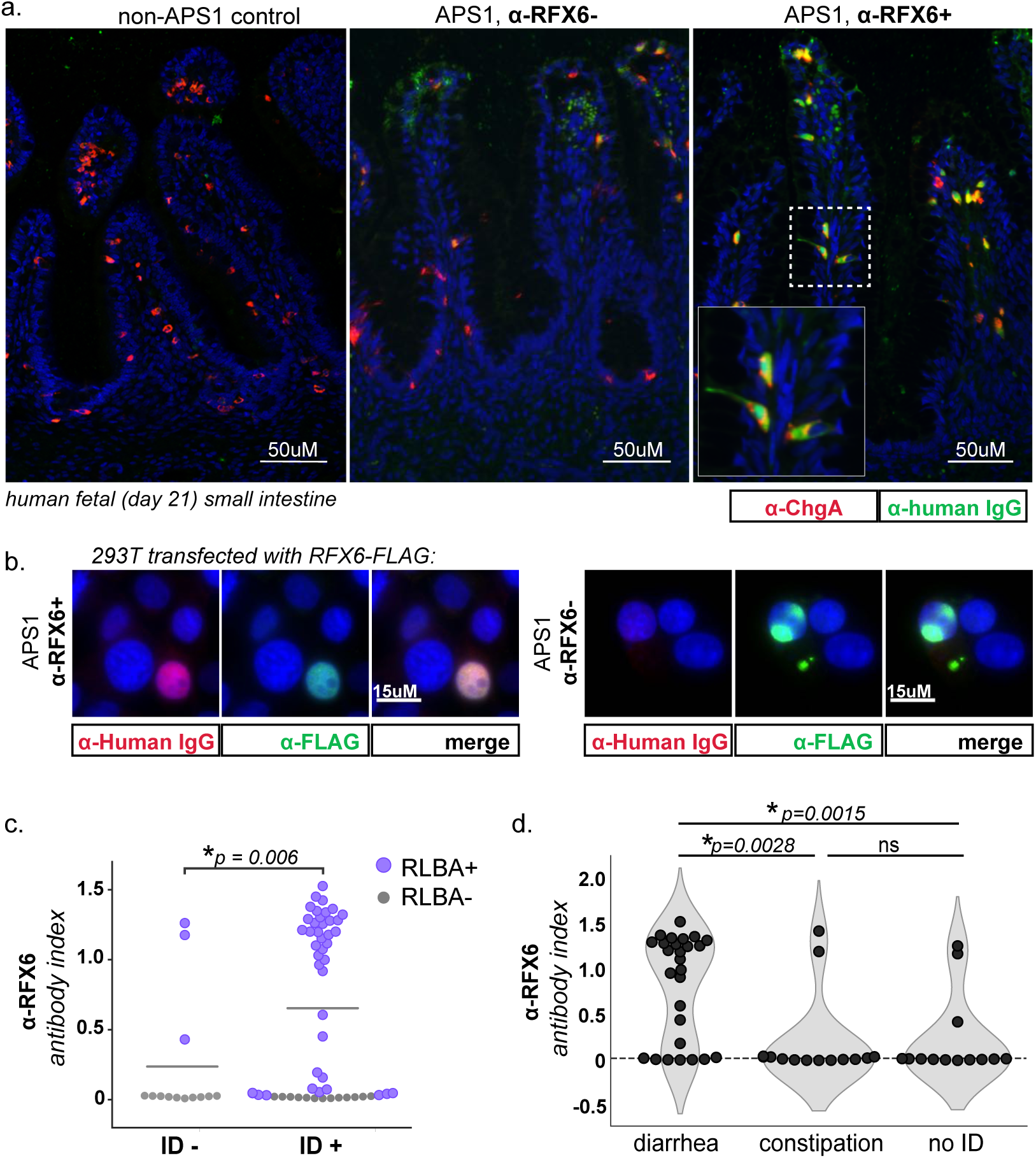
APS1 patients with intestinal dysfunction mount an antibody response to intestinal enteroendocrine cells and to enteroendocrine-expressed protein RFX6. **A.** Anti-RFX6 positive APS1 serum with intestinal dysfunction co-stains Chromogranin-A (ChgA) positive enteroendocrine cells in a nuclear pattern (right panel & inset). In contrast, non-APS1 control sera as well as anti-RFX6 negative APS1 serum do not co-stain ChgA+ enteroendocrine cells (left and center panels). **B.** Anti-RFX6+ serum, but not anti-RFX6-serum, co-stains HEK293T cells transfected with an RFX6-expressing plasmid. See also Figure 6: Figure Supplement 2. **C.** Radioligand binding assay (RLBA) anti-RFX6 antibody index is significantly higher across individuals with intestinal dysfunction (ID; Mann-Whitney U, p = 0.006). Purple color indicates samples that fall above 6 standard deviations of the mean non-APS1 control RLBA antibody index**. D.** Individuals with the diarrheal subtype of ID have a higher frequency of anti-RFX6 antibody positivity as compared to those with constipation-type ID (Mann-Whitney U, p=0.0028) or no ID (p=0.0015). See also Figure 6: Figure Supplement 3. For associated RFX6 PhIP-Seq & tissue expression data, see Figure 6: Figure Supplement 1.

Both mice and humans with biallelic mutation of the gene encoding RFX6 have enteroendocrine cell deficiency and intestinal malabsorption (Mitchell et al., 2004; Piccand et al., 2019; S. B. Smith et al., 2010), and humans with other forms of genetic or acquired enteroendocrine cell deficiency also suffer from chronic malabsorptive diarrhea (Akoury et al., 2015; Li et al., 2008; Reddy et al., 2012; X. Wang et al., 2018; W. Zhang et al., 2019). In this cohort, 54/67 (81%) of individuals have intestinal dysfunction defined as the presence of chronic diarrhea, chronic constipation or an alternating pattern of both, without meeting ROME III diagnostic criteria for irritable bowel syndrome, as previously described (Ferré et al., 2016). When the cohort was subsetted by presence or absence of intestinal dysfunction, the anti-RFX6 RLBA signal was significantly higher when intestinal dysfunction was present (**Figure 6C**). Further subsetting of the cohort by subtype of intestinal dysfunction revealed that individuals with anti-RFX6 antibodies belonged preferentially to the diarrheal-type (as opposed to constipation-type) group of intestinal dysfunction (**Figure 6D & Figure 6: Figure Supplement 3A**). Given that RFX6 is also expressed in the pancreas, we also examined the association of anti-RFX6 antibodies with APS1-associated type 1 diabetes. We observed that 6/7 APS1-associated type 1 diabetes samples had positive signal for anti-RFX6 antibodies by RLBA (**Figure 6: Figure Supplement 3B**). However, due to small sample size, an expanded cohort would be needed to determine the significance of this observation. Together, these data suggest that RFX6 is a common, shared autoantigen in APS1 that may be involved in the immune response to intestinal enteroendocrine cells as well as pancreatic islets. Future studies will help to determine whether testing for anti-RFX6 antibodies possesses clinical utility for prediction or diagnosis of specific APS1 autoimmune disease manifestations as well as for non-APS1 autoimmune disease.

## DISCUSSION

Here, we have identified a new set of autoantigens that are associated with autoimmune features in APS1 by using the broad-based antigen screening platform of PhIP-Seq. Unlike fixed protein arrays, programmable phage display possesses the advantage of being able to comprehensively cover all annotated proteins and their isoforms. The PhIP-Seq library used here is composed of over 700,000 peptides, each 49 amino acids, and corresponding to approximately 20,000 proteins and their known splicing isoforms. This is highly complementary to recently published protein arrays that cover approximately 9,000 distinct proteins (Fishman et al., 2017; Landegren et al., 2016; Meyer et al., 2016). Recent protein array approaches with APS1 samples using strict cutoffs have been able to identify a number of new autoantigen targets that include PDILT and MAGEB2 (Landegren et al., 2016). Several new targets, including RFX6, KHDC3L, ACP4, NKX6-3, ASMT, and PDX1, were likely discovered here because these antigens were not present on previously published protein array platforms. Only a subset of the novel targets identified here were validated orthogonally. While none failed validation relative to non-APS1 controls, further validation work will be needed for the many additional novel targets identified by PhIP-Seq. It is also worth mentioning that the PhIP-Seq method leverages continuing declines in the cost of oligonucleotide synthesis and Next-Generation Sequencing. Both technologies benefit from economies of scale, and once constructed, a PhIP-Seq phage library may be propagated in large quantities at negligible cost. The primary disadvantage of PhIP-Seq is the fact that conformation specific antibodies are likely to be missed, unless short linear subsequences carry significant binding energy. For example, PhIP-Seq detected only limited signal towards some literature reported antigens, including GAD65 and interferon family proteins in this APS1 cohort. Given that these antigens have been reported to involve conformational epitopes, antibodies to these antigens would not be predicted to be easily detected by linear peptides (Björk et al., 1994; Meager et al., 2006; Meyer et al., 2016; Wolff et al., 2013; Ziegler et al., 1996). Nonetheless, the ability to detect anti-interferon antibodies in a subset of APS1 samples highlights the utility of PhIP-Seq for antigen discovery despite decreased sensitivity for certain epitopes (**Figure 1: Figure Supplement 1**).

People with (Anderson, 2002; Cheng & Anderson, 2018; Husebye et al., 2018; Malchow et al., 2016)APS1 develop autoimmune manifestations over the course of many years, and it is thought that each manifestation may be explained by autoimmune response to one or few initial protein targets. In principle, these target proteins would most likely (1) exhibit thymic AIRE-dependency and (2) be restricted to the single or narrow range of tissues associated with the corresponding autoimmune disease. For example, adrenal insufficiency, which results from autoimmune response to cells of the adrenal gland, is thought to occur due to targeting of adrenally-expressed cytochrome p450 family members (Obermayer-Straub et al., 2000; Winqvist et al., 1993). However, a more complete understanding of the protein target spectrum paired with clinical phenotypic associations has been lacking. This, combined with the limited applicability of murine observations to the human disease, has left the question of which clinical characteristics best associate with APS1 autoantigens a heavily debated subject (Pöntynen et al., 2006).

Testing for defined autoantibody specificities provides substantial clinical benefit for prediction and diagnosis of autoimmune disease. A primary goal of this study was to identify autoantigens with potential clinical significance; consistently, our analyses focused primarily on antigens that appeared across multiple samples, rather than autoantigens that were restricted to individual samples. Using conservative inclusion criteria, we discovered 72 novel autoantigens that were shared across a minimum of 3 APS1 samples, of which 7/7 were successfully validated at the whole protein level. Overall, we have expanded the known repertoire of common APS1 antigens, confirming that the antibody target repertoire of common antigens in APS1 is larger than previously appreciated. Interestingly, our data also suggest that the size of the commonly autoantibody-targeted repertoire of proteins is dramatically lower than the number of genes (∼4000) that exhibit AIRE-dependent thymic expression.

The spectrum of different autoimmune diseases that can be observed in APS1 is extensive and has continued to expand through investigation of larger cohorts (Ahonen et al., 1990; Bruserud et al., 2016; Ferré et al., 2016). In this study, clinical metadata encompassing disease status across 24 individual disease manifestations in a total of 67 people with APS1 was leveraged to uncover (among others) an association of anti-KHDC3L antibodies and ovarian insufficiency, a disease that affects over half of all women with APS1 and manifests as abnormal menstrual cycling, reduced fertility, and early menopause. While autoreactivity to the steroidogenic granulosa cells – the cells surrounding and supporting the oocytes – has been proposed as one etiology of the clinical ovarian insufficiency, it has also been suggested that there may exist an autoimmune response to the oocyte itself (Jasti et al., 2012; Maclaren et al., 2001; Obermayer-Straub et al., 2000; Otsuka et al., 2011; Welt, 2008). Our finding that females with APS1-associated ovarian insufficiency exhibit autoantibodies to KHDC3L, an oocyte specific protein, supports this hypothesis. As exemplified by autoantibody presence in other autoimmune conditions, anti-KHDC3L antibodies may also have predictive value. Specifically, in our cohort, we found anti-KHDC3L antibodies to be present in many young, pre-menstrual females; these observations will require additional studies in prospective, longitudinal cohorts for further evaluation of potential predictive value. Interestingly, primary ovarian insufficiency (POI) in the absence of AIRE-deficiency is increasingly common and affects an estimated 1 in 100 women; up to half of these cases have been proposed to have autoimmune etiology (Huhtaniemi et al., 2018; Jasti et al., 2012; Nelson, 2009; Silva et al., 2014).

We noted that the majority of samples with antibodies to KHDC3L also exhibited antibodies to NLRP5, and vice versa. Remarkably, both of these proteins are critical parts of a subcortical maternal complex (SCMC) in both human and murine oocytes (Li et al., 2008; K. Zhu et al., 2015). Indeed, “multi-pronged” targeting of the same pathway has been previously implicated in APS1, where antibodies to DDC and TPH1 – enzymes in the serotonin and melatonin synthesis pathways – have been described (Ekwall et al., 1998; Husebye et al., 1997; Kluger et al., 2015). In addition to these targets, our data revealed an additional autoantibody-targeted enzyme ASMT in the same melatonin synthesis pathway. While the earlier TPH1- and DDC-catalyzed steps occur in both the intestine and pineal gland and precede the formation of serotonin, ASMT is predominantly expressed in the pineal gland and catalyzes the last, post-serotonin step in melatonin synthesis, suggesting that targeting of this pathway occurs at multiple distinct steps. To our knowledge, this is the first reported autoantigen in APS1 whose expression is restricted to the central nervous system.

In past and ongoing investigations, some individuals with APS1 have been reported to feature histologic loss of intestinal enteroendocrine cells on biopsy (Högenauer et al., 2001; Oliva-Hemker et al., 2006; Posovszky et al., 2012, Natarajan et al., manuscript in preparation). The association of anti-RFX6 antibodies with the diarrheal type of intestinal dysfunction is consistent with published studies in murine models of *Rfx6* (and enteroendocrine cell) ablation (Piccand et al., 2019; S. B. Smith et al., 2010). In addition, human enteroendocrine cell deficiency as well as mutations in enteroendocrine gene *NEUROG3* have been linked to chronic diarrhea and malabsorption, and recently, intestinal enteroendocrine cells have been suggested to play a role in mediating intestinal immune tolerance (Ohsie et al., 2009; Sifuentes-Dominguez et al., 2019; J. Wang et al., 2006). In sum, although APS1-associated intestinal dysfunction may have multiple etiologies, including autoimmune enteritis or dysfunction of exocrine pancreas, our findings of highly prevalent anti-RFX6 antibodies provide evidence of a common, shared autoantigen involved with this disease phenotype. In addition, patients with type 1 diabetes alone (not in association with APS1) frequently exhibit intestinal dysfunction related to multiple etiologies including Celiac disease, autonomic neuropathy, and exocrine pancreatic insufficiency (Du et al., 2018); future studies will be needed to determine whether anti-RFX6 antibodies may distinguish a subset of these patients with an autoimmune enteroendocrinopathy contributing to their symptoms.

While we report many novel antigens, we also acknowledge that the relationship between autoantibody status and disease is often complicated. This concept can be illustrated by examining the well-established autoantibody specificities in autoimmune diabetes (Taplin & Barker, 2009). First, islet autoantibodies (GAD65, ZNT8, etc.) can be found within non-autoimmune sera, where they are thought to represent an increased risk of developing disease as compared to the antibody-negative population. Second, not all patients with autoimmune diabetes are autoantibody positive. In sum, while autoantibodies can be extremely useful for risk assessment as well as for diagnosis, they often lack high sensitivity and specificity; both of these caveats can result in difficulties detecting strong clinical associations. For example, anti-ACP4 antibodies are highly prevalent in our cohort, but they exhibit only a trending association with dental enamel hypoplasia despite the strong biological evidence that ACP4 dysfunction leads to enamel hypoplasia (Seymen et al., 2016; C. E. Smith et al., 2017). Our data in humans is currently insufficient to determine whether immune responses to novel antigens such as ACP4 are pathogenic, indirectly linked to risk of disease, or instead simply represent a B-cell bystander effect. To better address these questions, we propose that future studies in mouse models could elucidate whether immune response to specific proteins, including ACP4, can result in the proposed phenotypes.

As the spectrum of diseases with potential autoimmune etiology continues to expand, the characteristic multiorgan autoimmunity in APS1 provides an ideal model system to more broadly approach the question of which proteins and cell types tend to be aberrantly targeted by the immune system. The data presented here has illuminated a collection of novel human APS1 autoimmune targets, as well as a novel antibody-disease association between RFX6 and diarrheal-type intestinal dysfunction, a highly prevalent disorder in APS1 that has until now lacked clinically applicable predictive or diagnostic markers. In sum, this data has significantly expanded the known autoantigen target profile in APS1 and highlighted several new directions for exploring the mechanics and clinical consequences of this complex syndrome.

## MATERIALS AND METHODS

### Data collection

All patient cohort data was collected and evaluated at the NIH, and all APECED/APS1 patients were enrolled in a research study protocols approved by the NIAID, NIH Clinical Center, and NCI Institutional Review Board Committee and provided with written informed consent for study participation. All NIH patients gave consent for passive use of their medical record for research purposes (protocol #11-I-0187). The majority of this cohort data was previously published by Ferré et al. 2016 and Ferré et al. 2019.

### Phage Immunoprecipitation – Sequencing (PhIP-Seq)

For PhIP-Seq, we adapted a custom-designed phage library consisting of 731,724 49AA peptides tiling the full protein-coding human genome including all isoforms (as of 2016) with 25AA overlap as previously described (O’Donovan et al., 2018). 1 milliliter of phage library was incubated with 1 microliter of human serum overnight at 4C, and human antibody (bound to phage) was immunoprecipitated using 40ul of a 1:1 mix of protein A/G magnetic beads (Thermo Fisher, Waltham, MA, #10008D & #10009D). Beads were washed 4 times and antibody-bound phage were eluted into 1ml of E. Coli at OD of 0.5-0.7 (BLT5403, EMD Millipore, Burlington, MA) for selective amplification of eluted phage. This library was re-incubated with human serum and repeated, followed by phenol-chloroform extraction of DNA from the final phage library. DNA was barcoded and amplified (Phusion PCR, 30 rounds), gel purified, and subjected to Next-Generation Sequencing on an Illumina MiSeq Instrument (Illumina, San Diego, CA).

### PhIP-Seq Analysis

Sequencing reads from fastq files were aligned to the reference oligonucleotide library and peptide counts were subsequently normalized by converting raw reads to percentage of total reads per sample. Peptide and gene-level enrichments for both APS1 and non-APS1 sera were calculated by determining the fold-change of read percentage per peptide and gene in each sample over the mean read percentage per peptide and gene in a background of mock-IP (A/G bead only, n = 18). Individual samples were considered positive for genes where the enrichment value was 10-fold or greater as compared to mock-IP. For plotting of multiple genes in parallel (**Figures 1 & 2**), enrichment values were z-scored and hierarchically clustered using Pearson correlation.

### Statistics

For comparison of distribution of PhIP-Seq gene enrichment between APS1 patients with and without specific disease manifestations, a (non-parametric) Kolmogorov-Smirnov test was used. For radioligand binding assays, antibody index for each sample was calculated as follows: (sample value – mean blank value) / (positive control antibody value – mean blank value). Comparison of antibody index values between non-APS1 control samples and APS1 samples was performed using a Mann-Whitney *U* test. Experimental samples that fell 3 standard deviations above of the mean of non-APS1 controls for each assay were considered positive, except in the case of RFX6, where a cutoff of 6 standard deviations above the mean of non-APS1 controls was used.

### Assessing tissue-specific RNA expression

To determine tissue-specificity and tissue-restriction of *Rfx6* expression in mice, we used publicly available Tabula Muris data (tabula-muris.ds.czbiohub.org) (Schaum et al., 2018). For investigation of *KHDC3L* expression in human ovary, we downloaded publicly available normalized FPKM transcriptome data from human oocytes and granulosa cells (GSE107746_Folliculogenesis_FPKM.log2.txt) (Y. Zhang et al., 2018). With this data, we performed principle component analysis, which clustered the two cell types correctly according to their corresponding sample label, and plotted log2(FPKM) by color for each sample.

### 293T overexpression assays

Human kidney embryo 293T (ATCC, Manassas, VA, #CRL-3216) cells were plated at 30% density in a standard 24-well glass bottom plate in complete DMEM media (Thermo Fisher, #119651198) with 10% Fetal Bovine Serum (Thermo Fisher, #10438026), 292ug/ml L-glutamine, 100ug/ml Streptomycin Sulfate, and 120Units/ml of Penicillin G Sodium (Thermo Fisher, #10378016). 18 hours later, cells were transiently transfected using a standard calcium chloride transfection protocol. For transfections, 0.1ug of sequence-verified pCMV-insert-MYC-FLAG overexpression vectors containing either no insert (Origene #PS100001; ‘mock’ transfection) or RFX6 insert (Origene #RC206174) were transfected into each well. 24 hours post-transfection, cells were washed in 1X PBS and fixed in 4% PFA for 10 minutes at room temperature.

### 293T indirect immunofluorescence

Fixed 293T cells were blocked for 1 hour at room temperature in 5% BSA in PBST. For primary antibody incubation, cells were incubated with human serum (1:1000) and rabbit anti-FLAG antibody (1:2000) in 5% BSA in PBST for 2 hours at room temperature (RT). Cells were washed 4X in PBST and subsequently incubated with secondary antibodies (goat anti-rabbit IgG 488, Invitrogen, Carlsbad, CA; #A-11034, 1:4000; & goat anti-human 647, Invitrogen #A-21445, 1:4000) for 1 hour at room temperature. Finally, cells were washed 4X in PBST, incubated with DAPI for 5 minutes at RT, and subsequently placed into PBS for immediate imaging. All images were acquired with a Nikon Ti inverted fluorescence microscope (Nikon Instruments, Melville, NY). All experiments were performed in biological duplicates.

### Indirect dual immunofluorescence on human fetal intestine

Human fetal small bowels (21.2 days gestational age) were processed as previously described (Berger et al., 2015). Individual APS1 sera (1:4000 dilution) were used in combination with rabbit antibodies to human Chromogranin A (Abcam, Cambridge, MA; #ab15160, 1:5000 dilution). Immunofluorescence detection utilized secondary Alexa Fluor secondary antibodies (Life Technologies, Waltham, MA; 488 goat anti-human IgG, #A11013; & 546 goat anti-rabbit IgG, #A11010). Nuclear DNA was stained with Hoechst dye (Invitrogen, #33342). All images were acquired with a Leica SP5 White Light confocal laser microscope (Leica Microsystems, Buffalo Grove, IL).

### 35S-radiolabeled protein generation and binding assay

DNA plasmids containing full-length cDNA under the control of a T7 promoter for each of the validated antigens (**Supplemental Table 3**) were verified by Sanger sequencing and used as DNA templates in the T7 TNT in vitro transcription/translation kit (Promega, Madison, WI; #L1170) using [35S]-methionine (PerkinElmer, Waltham, MA; #NEG709A). Protein was column-purified on Nap-5 columns (GE healthcare, Chicago, IL; #17-0853-01) and immunoprecipitated on Sephadex protein A/G beads (Sigma Aldrich, St. Louis, MO; #GE17-5280-02 and #GE17-0618-05, 4:1 ratio) in duplicate with serum or control antibodies in 96-well polyvinylidene difluoride filtration plates (Corning, Corning, NY; #EK-680860). Each well contained 35’000 counts per minute (cpm) of radiolabeled protein and 2.5ul of serum or appropriately diluted control antibody (**Supplemental Table 3**). The cpms of immunoprecipitated protein was quantified using a 96-well Microbeta Trilux liquid scintillation plate reader (Perkin Elmer).

## ACKNOWLEDGEMENTS

We thank Joseph M. Replogle, Jeffrey A. Hussmann, Madhura Raghavan, Hanna Retallack, Brian D. O’Donovan, and members of the DeRisi, Anderson, Lionakis, and German labs for helpful discussions. We thank Kari Herrington and the UCSF Nikon Imaging Center for imaging support, as well as Sabrina Mann, Wint Lwin, and the UCSF Center for Advanced Technology for technical support. We thank the New York Blood Bank for providing us with the de-identified human non-inflammatory control plasma samples used in this study.

## SUPPLEMENTAL FIGURES

**Supplemental Table 1.**
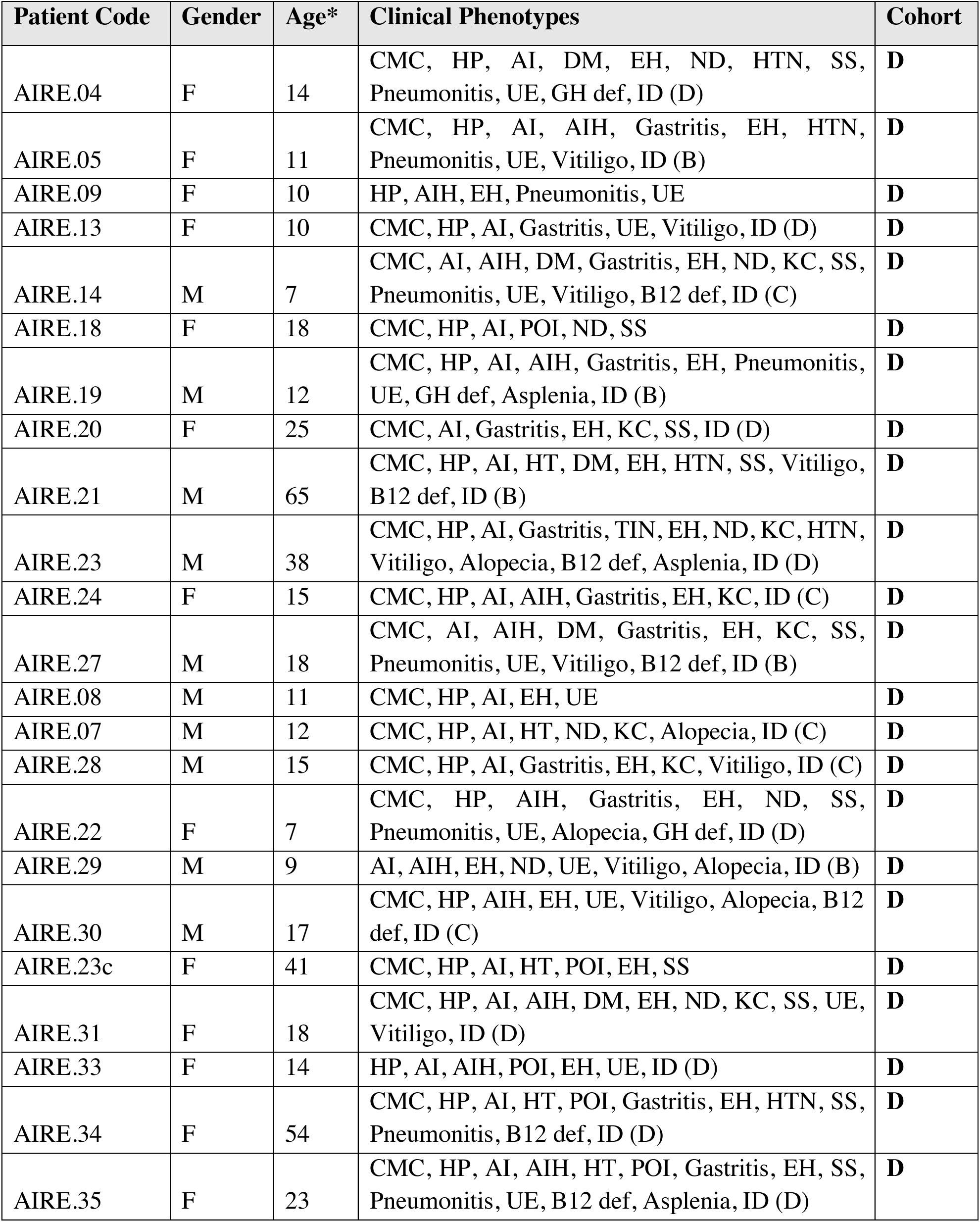
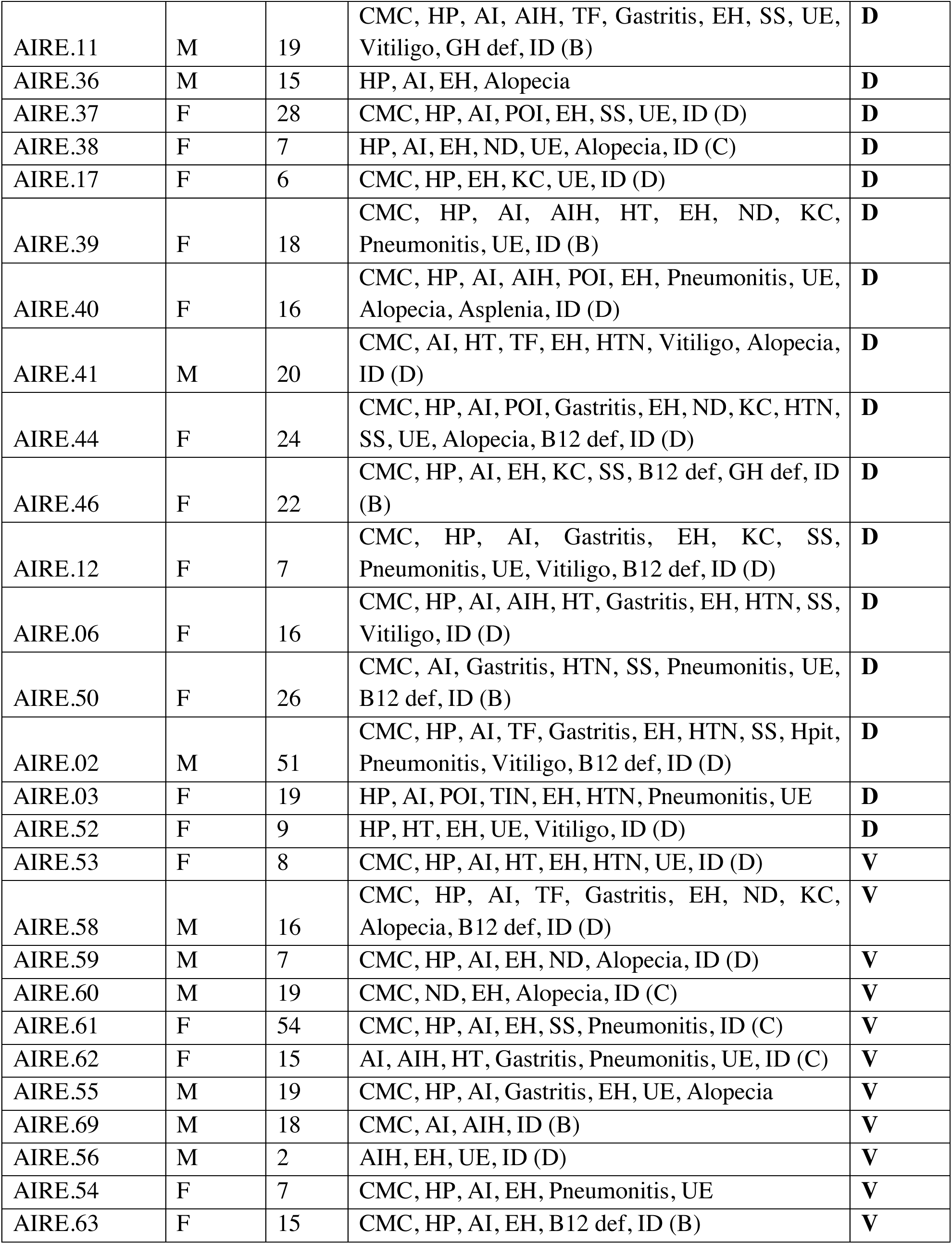
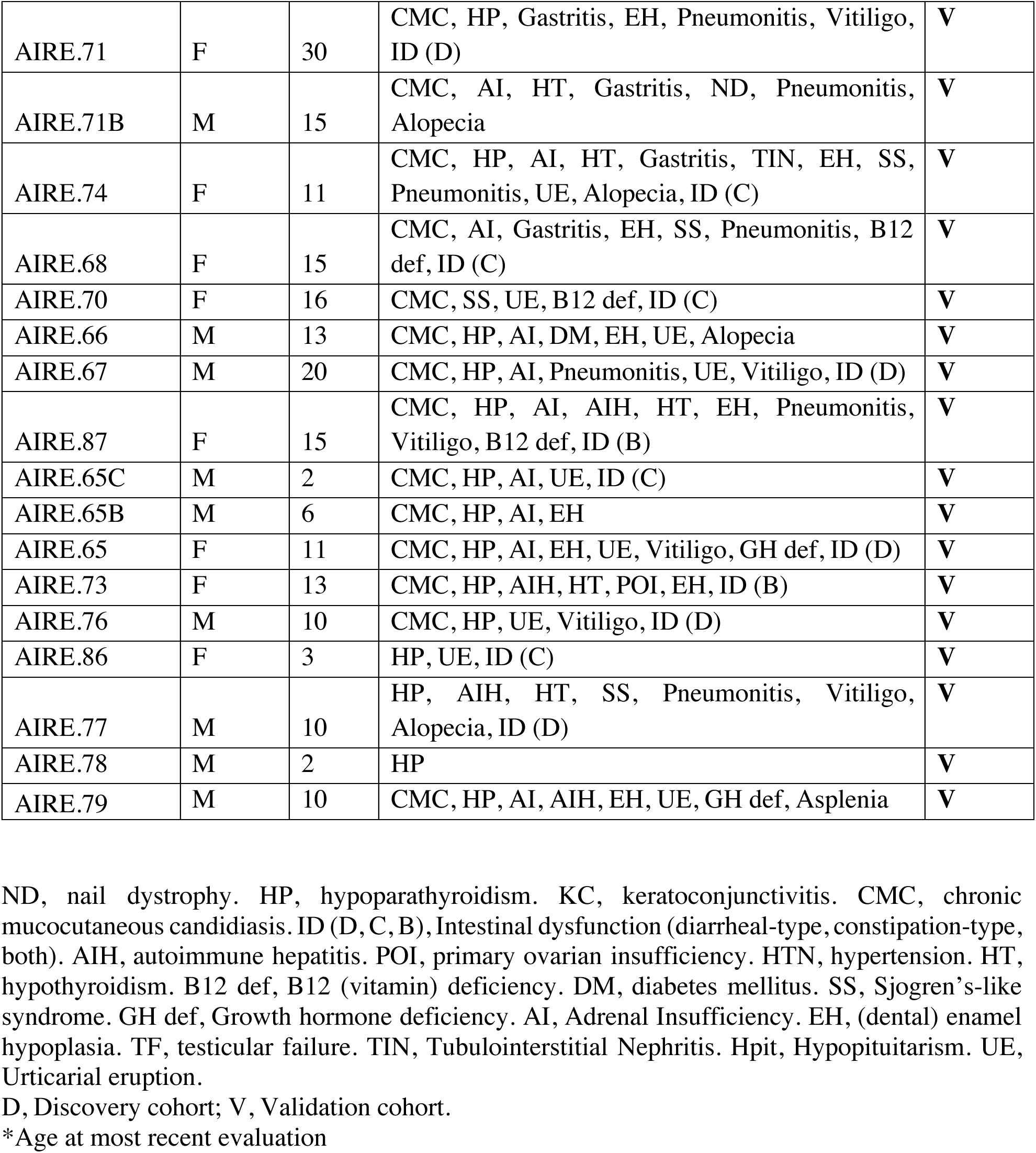
APS1 cohort: Clinical Data.

| Patient Code | Gender | Age* | Clinical Phenotypes | Cohort |
| --- | --- | --- | --- | --- |
| AIRE.04 | F | 14 | CMC, HP, AI, DM, EH, ND, HTN, SS, Pneumonitis, UE, GH def, ID (D) | **D** |
| AIRE.05 | F | 11 | CMC, HP, AI, AIH, Gastritis, EH, HTN, Pneumonitis, UE, Vitiligo, ID (B) | **D** |
| AIRE.09 | F | 10 | HP, AIH, EH, Pneumonitis, UE | **D** |
| AIRE.13 | F | 10 | CMC, HP, AI, Gastritis, UE, Vitiligo, ID (D) | **D** |
| AIRE.14 | M | 7 | CMC, AI, AIH, DM, Gastritis, EH, ND, KC, SS, Pneumonitis, UE, Vitiligo, B12 def, ID (C) | **D** |
| AIRE.18 | F | 18 | CMC, HP, AI, POI, ND, SS | **D** |
| AIRE.19 | M | 12 | CMC, HP, AI, AIH, Gastritis, EH, Pneumonitis, UE, GH def, Asplenia, ID (B) | **D** |
| AIRE.20 | F | 25 | CMC, AI, Gastritis, EH, KC, SS, ID (D) | **D** |
| AIRE.21 | M | 65 | CMC, HP, AI, HT, DM, EH, HTN, SS, Vitiligo, B12 def, ID (B) | **D** |
| AIRE.23 | M | 38 | CMC, HP, AI, Gastritis, TIN, EH, ND, KC, HTN, Vitiligo, Alopecia, B12 def, Asplenia, ID (D) | **D** |
| AIRE.24 | F | 15 | CMC, HP, AI, AIH, Gastritis, EH, KC, ID (C) | **D** |
| AIRE.27 | M | 18 | CMC, AI, AIH, DM, Gastritis, EH, KC, SS, Pneumonitis, UE, Vitiligo, B12 def, ID (B) | **D** |
| AIRE.08 | M | 11 | CMC, HP, AI, EH, UE | **D** |
| AIRE.07 | M | 12 | CMC, HP, AI, HT, ND, KC, Alopecia, ID (C) | **D** |
| AIRE.28 | M | 15 | CMC, HP, AI, Gastritis, EH, KC, Vitiligo, ID (C) | **D** |
| AIRE.22 | F | 7 | CMC, HP, AIH, Gastritis, EH, ND, SS, Pneumonitis, UE, Alopecia, GH def, ID (D) | **D** |
| AIRE.29 | M | 9 | AI, AIH, EH, ND, UE, Vitiligo, Alopecia, ID (B) | **D** |
| AIRE.30 | M | 17 | CMC, HP, AIH, EH, UE, Vitiligo, Alopecia, B12 def, ID (C) | **D** |
| AIRE.23c | F | 41 | CMC, HP, AI, HT, POI, EH, SS | **D** |
| AIRE.31 | F | 18 | CMC, HP, AI, AIH, DM, EH, ND, KC, SS, UE, Vitiligo, ID (D) | **D** |
| AIRE.33 | F | 14 | HP, AI, AIH, POI, EH, UE, ID (D) | **D** |
| AIRE.34 | F | 54 | CMC, HP, AI, HT, POI, Gastritis, EH, HTN, SS, Pneumonitis, B12 def, ID (D) | **D** |
| AIRE.35 | F | 23 | CMC, HP, AI, AIH, HT, POI, Gastritis, EH, SS, Pneumonitis, UE, B12 def, Asplenia, ID (D) | **D** |
| AIRE.11 | M | 19 | CMC, HP, AI, AIH, TF, Gastritis, EH, SS, UE, Vitiligo, GH def, ID (B) | **D** |
| AIRE.36 | M | 15 | HP, AI, EH, Alopecia | **D** |
| AIRE.37 | F | 28 | CMC, HP, AI, POI, EH, SS, UE, ID (D) | **D** |
| AIRE.38 | F | 7 | HP, AI, EH, ND, UE, Alopecia, ID (C) | **D** |
| AIRE.17 | F | 6 | CMC, HP, EH, KC, UE, ID (D) | **D** |
| AIRE.39 | F | 18 | CMC, HP, AI, AIH, HT, EH, ND, KC, Pneumonitis, UE, ID (B) | **D** |
| AIRE.40 | F | 16 | CMC, HP, AI, AIH, POI, EH, Pneumonitis, UE, Alopecia, Asplenia, ID (D) | **D** |
| AIRE.41 | M | 20 | CMC, AI, HT, TF, EH, HTN, Vitiligo, Alopecia, ID (D) | **D** |
| AIRE.44 | F | 24 | CMC, HP, AI, POI, Gastritis, EH, ND, KC, HTN, SS, UE, Alopecia, B12 def, ID (D) | **D** |
| AIRE.46 | F | 22 | CMC, HP, AI, EH, KC, SS, B12 def, GH def, ID (B) | **D** |
| AIRE.12 | F | 7 | CMC, HP, AI, Gastritis, EH, KC, SS, Pneumonitis, UE, Vitiligo, B12 def, ID (D) | **D** |
| AIRE.06 | F | 16 | CMC, HP, AI, AIH, HT, Gastritis, EH, HTN, SS, Vitiligo, ID (D) | **D** |
| AIRE.50 | F | 26 | CMC, AI, Gastritis, HTN, SS, Pneumonitis, UE, B12 def, ID (B) | **D** |
| AIRE.02 | M | 51 | CMC, HP, AI, TF, Gastritis, EH, HTN, SS, Hpit, Pneumonitis, Vitiligo, B12 def, ID (D) | **D** |
| AIRE.03 | F | 19 | HP, AI, POI, TIN, EH, HTN, Pneumonitis, UE | **D** |
| AIRE.52 | F | 9 | HP, HT, EH, UE, Vitiligo, ID (D) | **D** |
| AIRE.53 | F | 8 | CMC, HP, AI, HT, EH, HTN, UE, ID (D) | **V** |
| AIRE.58 | M | 16 | CMC, HP, AI, TF, Gastritis, EH, ND, KC, Alopecia, B12 def, ID (D) | **V** |
| AIRE.59 | M | 7 | CMC, HP, AI, EH, ND, Alopecia, ID (D) | **V** |
| AIRE.60 | M | 19 | CMC, ND, EH, Alopecia, ID (C) | **V** |
| AIRE.61 | F | 54 | CMC, HP, AI, EH, SS, Pneumonitis, ID (C) | **V** |
| AIRE.62 | F | 15 | AI, AIH, HT, Gastritis, Pneumonitis, UE, ID (C) | **V** |
| AIRE.55 | M | 19 | CMC, HP, AI, Gastritis, EH, UE, Alopecia | **V** |
| AIRE.69 | M | 18 | CMC, AI, AIH, ID (B) | **V** |
| AIRE.56 | M | 2 | AIH, EH, UE, ID (D) | **V** |
| AIRE.54 | F | 7 | CMC, HP, AI, EH, Pneumonitis, UE | **V** |
| AIRE.63 | F | 15 | CMC, HP, AI, EH, B12 def, ID (B) | **V** |
| AIRE.71 | F | 30 | CMC, HP, Gastritis, EH, Pneumonitis, Vitiligo, ID (D) | V |
| AIRE.71B | M | 15 | CMC, AI, HT, Gastritis, ND, Pneumonitis, Alopecia | V |
| AIRE.74 | F | 11 | CMC, HP, AI, HT, Gastritis, TIN, EH, SS, Pneumonitis, UE, Alopecia, ID (C) | V |
| AIRE.68 | F | 15 | CMC, AI, Gastritis, EH, SS, Pneumonitis, B12 def, ID (C) | V |
| AIRE.70 | F | 16 | CMC, SS, UE, B12 def, ID (C) | V |
| AIRE.66 | M | 13 | CMC, HP, AI, DM, EH, UE, Alopecia | V |
| AIRE.67 | M | 20 | CMC, HP, AI, Pneumonitis, UE, Vitiligo, ID (D) | V |
| AIRE.87 | F | 15 | CMC, HP, AI, AIH, HT, EH, Pneumonitis, Vitiligo, B12 def, ID (B) | V |
| AIRE.65C | M | 2 | CMC, HP, AI, UE, ID (C) | V |
| AIRE.65B | M | 6 | CMC, HP, AI, EH | V |
| AIRE.65 | F | 11 | CMC, HP, AI, EH, UE, Vitiligo, GH def, ID (D) | V |
| AIRE.73 | F | 13 | CMC, HP, AIH, HT, POI, EH, ID (B) | V |
| AIRE.76 | M | 10 | CMC, HP, UE, Vitiligo, ID (D) | V |
| AIRE.86 | F | 3 | HP, UE, ID (C) | V |
| AIRE.77 | M | 10 | HP, AIH, HT, SS, Pneumonitis, Vitiligo, Alopecia, ID (D) | V |
| AIRE.78 | M | 2 | HP | V |
| AIRE.79 | M | 10 | CMC, HP, AI, AIH, EH, UE, GH def, Asplenia | V |

ND, nail dystrophy. HP, hypoparathyroidism. KC, keratoconjunctivitis. CMC, chronic mucocutaneous candidiasis. ID (D, C, B), Intestinal dysfunction (diarrheal-type, constipation-type, both). AIH, autoimmune hepatitis. POI, primary ovarian insufficiency. HTN, hypertension. HT, hypothyroidism. B12 def, B12 (vitamin) deficiency. DM, diabetes mellitus. SS, Sjogren's-like syndrome. GH def, Growth hormone deficiency. AI, Adrenal Insufficiency. EH, (dental) enamel hypoplasia. TF, testicular failure. TIN, Tubulointerstitial Nephritis. Hpit, Hypopituitarism. UE, Urticarial eruption.
D, Discovery cohort; V, Validation cohort.
\*Age at most recent evaluation

**Supplemental Table 2.** Tissue-restricted expression patterns of validated novel APS1 antigens.

| Gene<br>(Human/mouse) | Protein Atlas: RNA<br>specificity category<br>(Tissue)<sup>14</sup> | Selected literature annotations |
| --- | --- | --- |
| *RFX6/Rfx6* | Tissue enhanced | Pancreas<br>Islets (Piccand et al., 2014; S. B. Smith et al., 2010)<br>Intestine<br>Enteroendocrine cells (Gehart et al., 2019; Piccand et al., 2019; S. B. Smith et al., 2010) |
| *KHDC3L/Khdc3* | Group enriched | Ovary<br>Oocytes (Y. Zhang et al., 2018; K. Zhu et al., 2015) |
| *ACP4/Acp4* | Tissue enhanced | Testes (Yousef et al., 2001)<br>Dental enamel (Green et al., 2019; Seymen et al., 2016; C. E. Smith et al., 2017) |
| *ASMT/Asmt* | Tissue enhanced | Brain<br>Pineal Gland (Rath et al., 2016) |
| *GIP/Gip* | Tissue enriched | Intestine<br>Enteroendocrine cells (Moody et al., 1984) |
| *NKX6-3 / Nkx6-3* | Group enriched | Pancreas<br>PP-cells (Schaum et al., 2018)<br>Intestine (Alanentalo et al., 2006) |
| *PDX1 / Pdx1* | Group enriched | Pancreas<br>Islets (Holland et al., 2002; Stoffers et al., 1997) |

**Supplemental Table 3.** Antibody information by application.

| antibody | Application<br>(IF:<br>immunofluorescence;<br>RLBA: radioligand<br>binding assay; CBA:<br>cell-based assay) | dilution |
| --- | --- | --- |
| Anti-NLRP5 (Santa Cruz, Dallas, TX; #sc-50630) | NLRP5 RLBA | 1:50 |
| Anti-SOX10 (Abcam, Cambridge, MA, #ab181466) | SOX10 RLBA | 1:25 |
| Anti-RFX6 (R&D Systems, Minneapolis, MN; #AF7780) | RFX6 RLBA | 1:50 |
| Anti-KHDC3L (Abcam, #ab170298) | KHDC3L RLBA | 1:25 |
| Anti-CYP11A1 (Abcam, #ab175408) | CYP11A1 RLBA | 1:50 |
| Anti-NKX6-3 (Biorbyt, Cambridge, Cambridgeshire, UK; #orb127108) | NKX6-3 RLBA | 1:50 |
| Anti-GIP (Abcam, #ab30679) | GIP RLBA | 1:50 |
| Anti-PDX1 (Invitrogen, Carlsbad, CA, #PA5-78024) | PDX1 RLBA | 1:50 |
| Anti-ASMT (Invitrogen, #PA5-24721) | ASMT RLBA | 1:25 |
| Anti-CHGA (Abcam, Cambridge, MA, USA, #ab15160) | Tissue IF | 1:5000 |
| Human serum | Tissue IF<br>CBA IF<br>RLBA | 1:4000<br>(Tissue)<br>1:500 (CBA)<br>1:25 (RLBA) |
| **Secondary abs:**<br>488 goat anti-human IgG (Life Technologies, Waltham, MA, USA; #A11013)<br>546 goat anti-rabbit IgG (Life Technologies, A11010) | Tissue IF | 1:400 |
| **Secondary abs:**<br>647 goat anti-human IgG (Thermo Fisher, #A-21445)<br>488 goat anti-rabbit IgG (Thermo Fisher, #A-11034) | CBA IF | 1:1000 |
| Anti-DYKDDDDK (D6W5B) (Cell Signaling Technologies, Danvers, MA; #14793) | CBA IF;<br>ACP4 RLBA | 1:2000 (CBA IF)<br>1:125 (RLBA) |
| **Nuclear staining:**<br>Hoechst dye (Invitrogen, #33342)<br>DAPI (Thermo Fisher, #D1306) | Tissue IF<br>CBA IF |  |

## FIGURE SUPPLEMENTS

**Figure 1: Figure Supplement 1.**
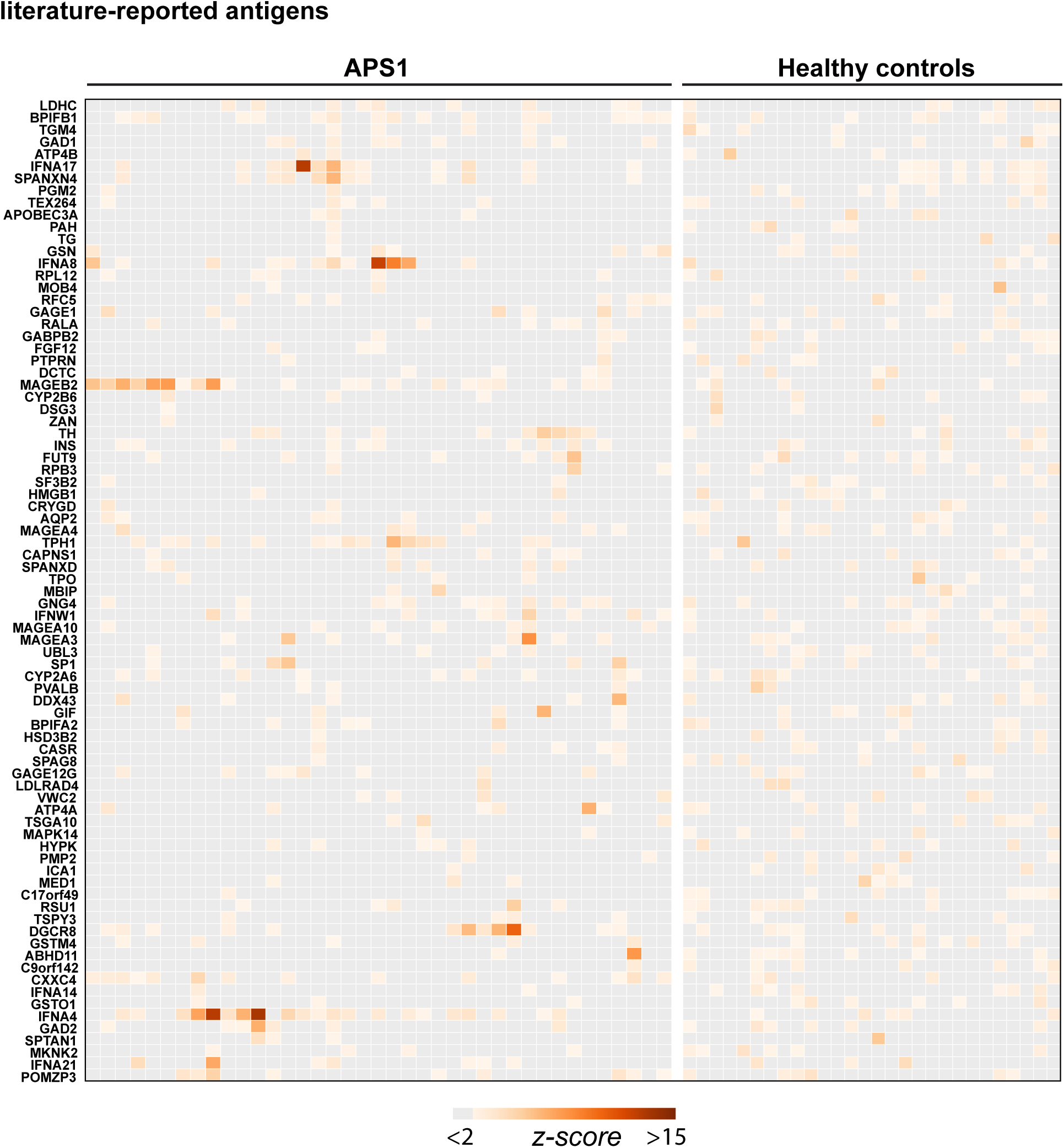
Hierarchically clustered (Pearson) Z-scored heatmap of literature reported autoantigens that did not meet the cutoff of 10-fold or greater signal over mock-IP in at least 2/39 APS1 sera and in 0/28 non-APS1 control sera.

**Figure 1: Figure Supplement 2.**
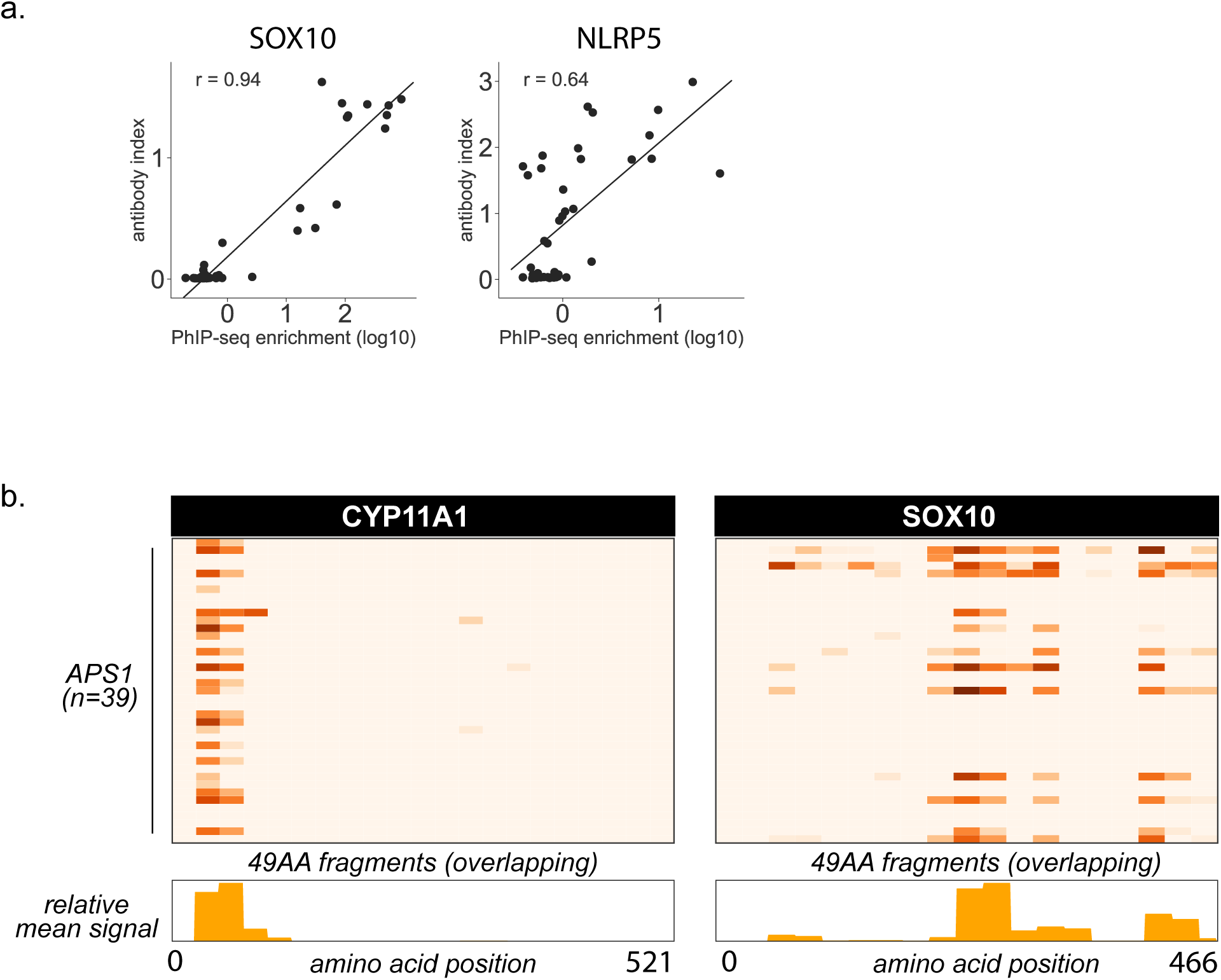
**A.** Scatterplot of individual PhIP-Seq enrichment values (log10) over mock-IP as compared to radioligand binding assay antibody index values (1 = commercial antibody signal) for known antigens SOX10 and NLRP5, with Pearson correlation coefficient r. **B.** PhIP-Seq enables 49 amino acid resolution of antibody signal from APS1 sera to literature-reported antigens CYP11A1 and SOX10. Top panels: PhIP-Seq signal (fold-change of each peptide as compared to signal from mock-IP, log10-scaled) for fragments 1-21 for CYP11A1 and fragments 1-19 for SOX10. Bottom panels: Trace of normalized signal for CYP11A1 and SOX10 fragments across the mean of all 39 APS1 sera.

**Figure 2: Figure Supplement 1.**
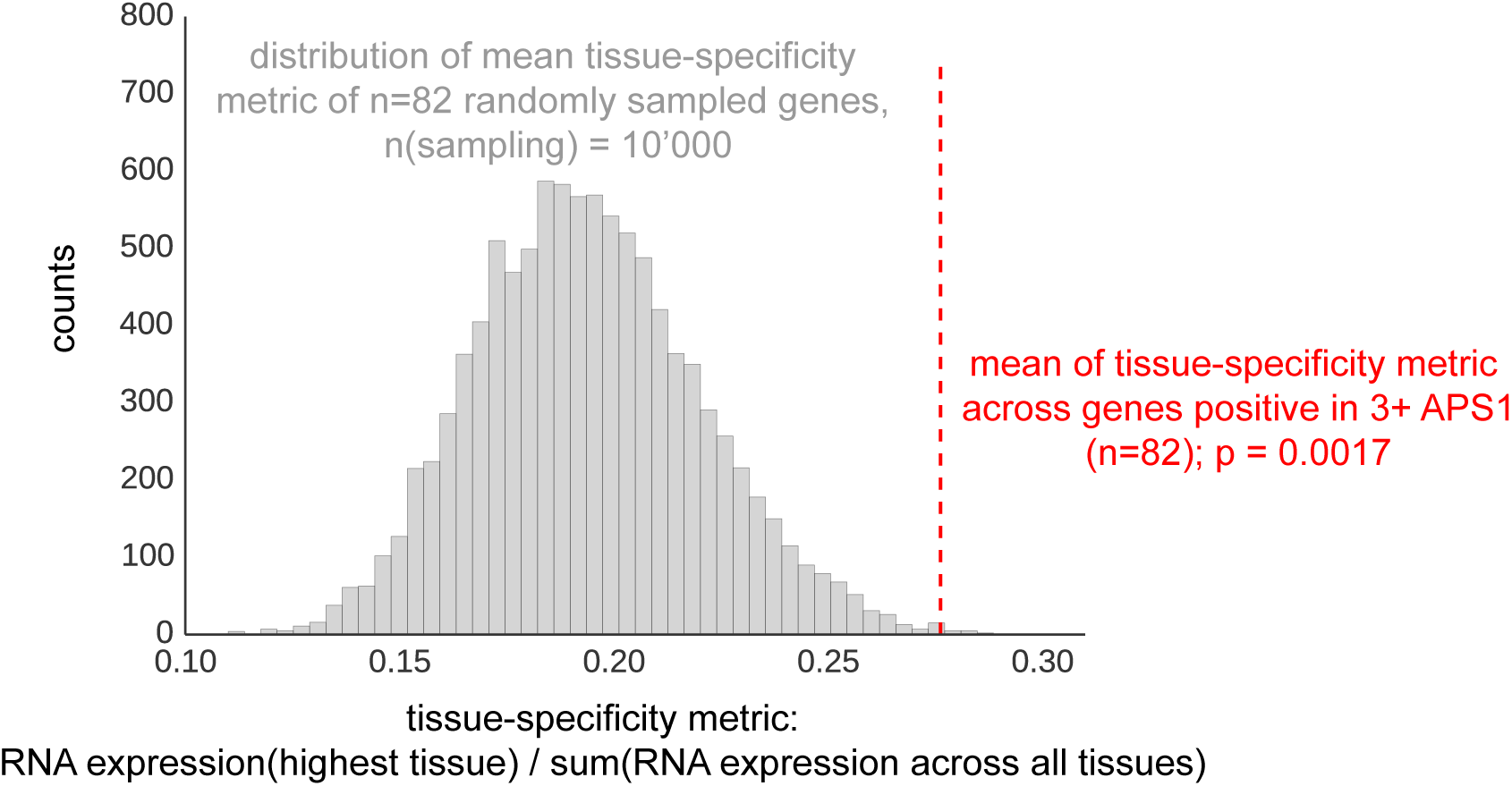
The mean of tissue-specificity ratio of 82 PhIP-Seq antigens (Figure 2) is increased as compared to the tissue-specificity ratio of n=82 randomly sampled genes (n-sampling = 10’000). Data from Protein Atlas, HPA/Gtex/Fantom5 RNA consensus dataset (Uhlen et al., 2015).

**Figure 3: Figure Supplement 1.**
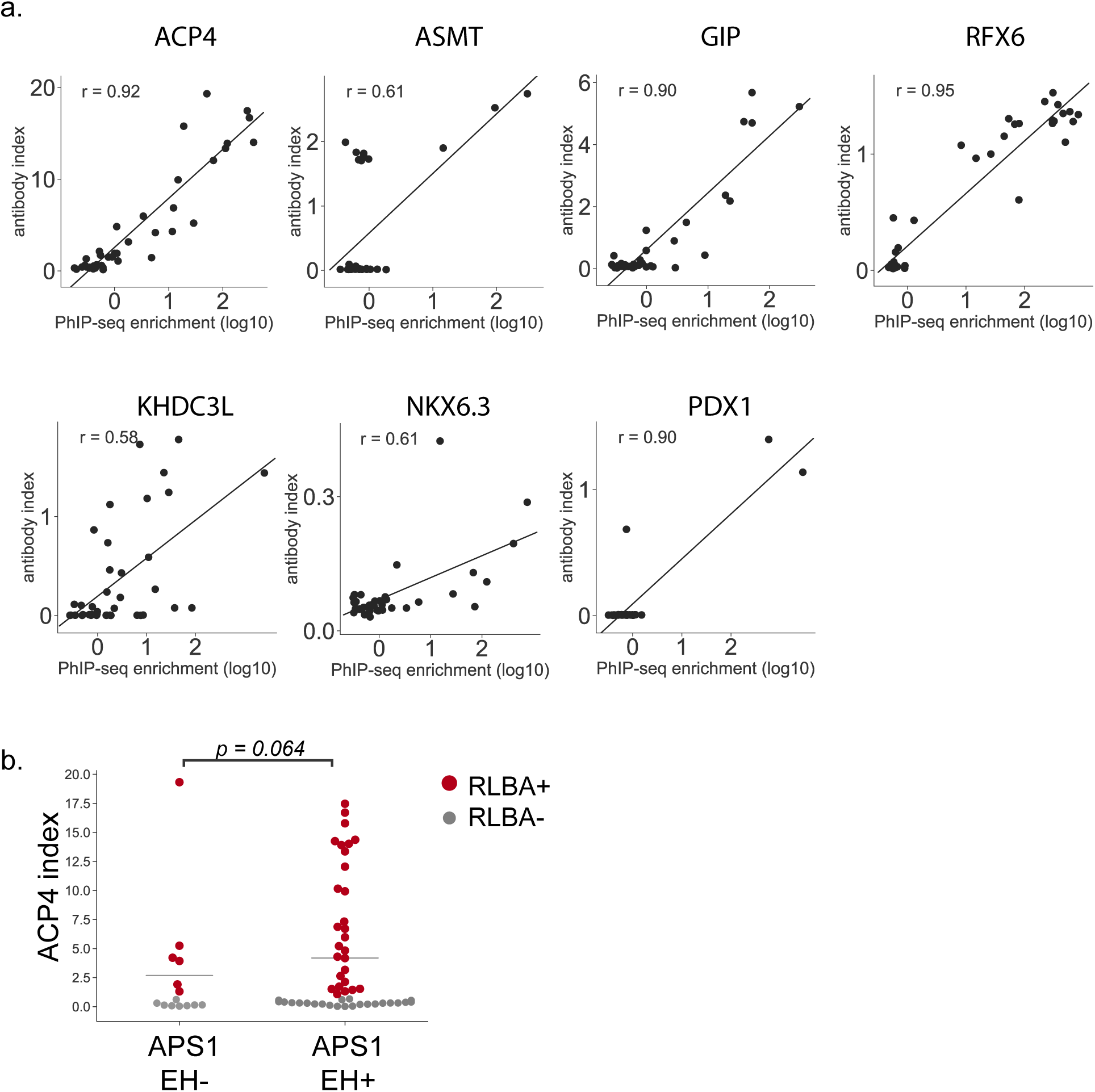
**A**. Scatterplot of individual PhIP-Seq enrichment values (log10) over mock-IP as compared to radioligand binding assay antibody index values (1 = commercial antibody signal) for novel antigens ACP4, ASMT, GIP, RFX6, KHDC3L, NKX6.3, and PDX1, with Pearson correlation coefficient r (Note that for PDX1, there are insufficient positive data points for the correlation to be meaningful). **B**. ACP4 RLBA autoantibody index, broken down by enamel hypoplasia (EH) status.

**Figure 4: Figure Supplement 1.**
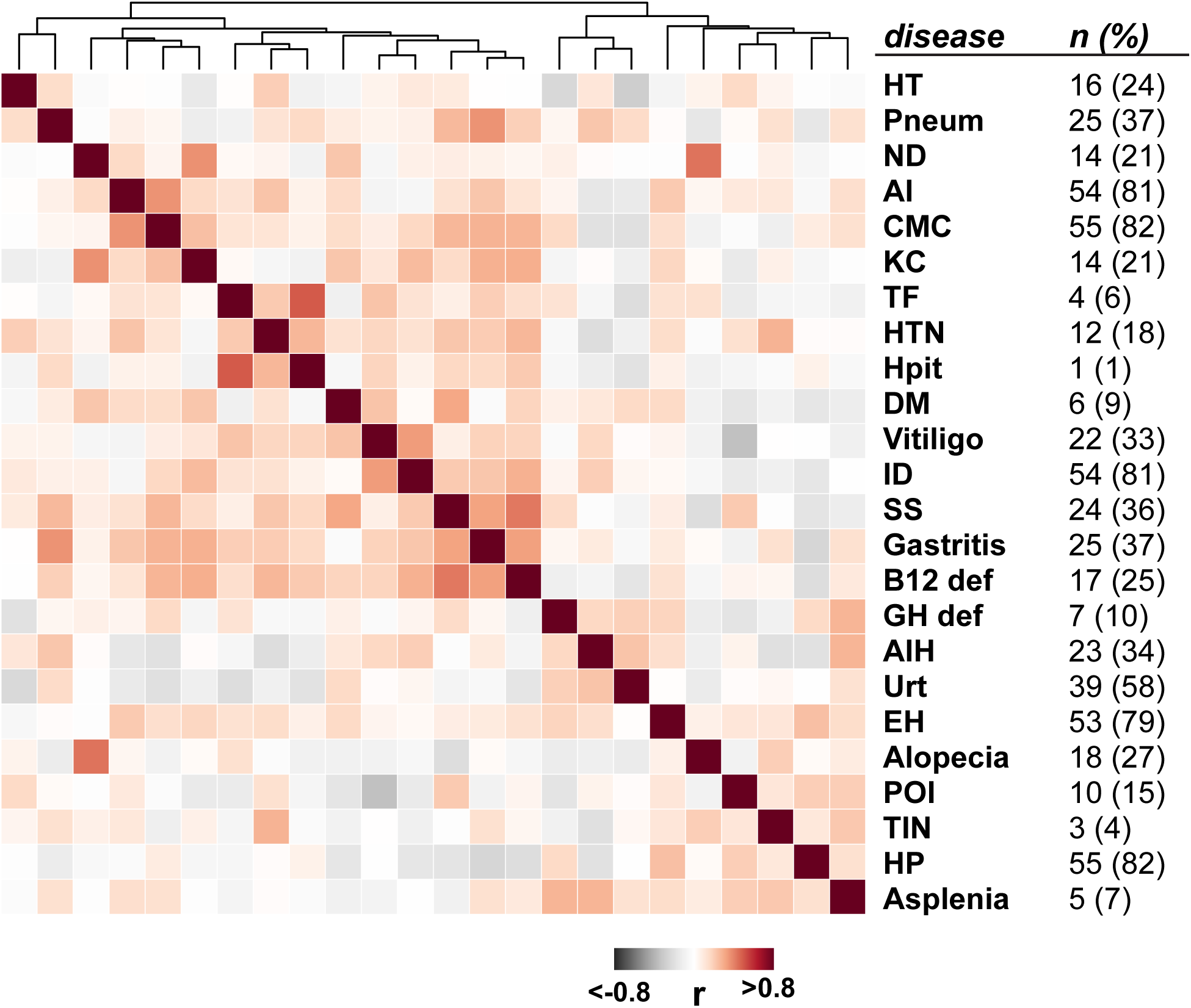
Clustered disease correlations in the APS1 cohort (Spearman’s rank correlation; n = 67).

**Figure 4: Figure Supplement 2.**
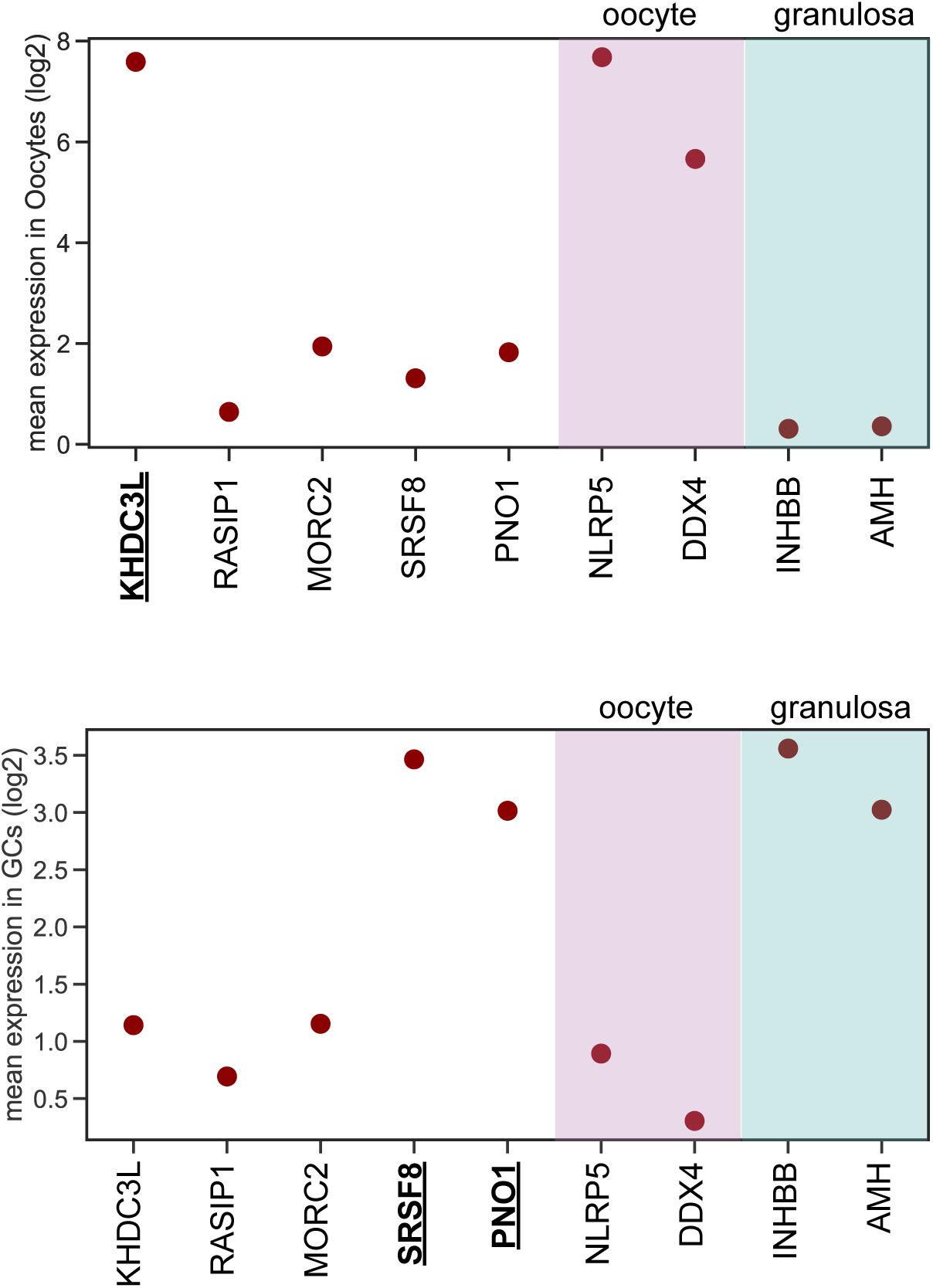
KHDC3L is highly expressed in oocytes (top), but not in granulosa cells (bottom). In contrast, SRSF8 and PNO1 are highly expressed in granulosa cells, but not in oocytes. Data from (Y. Zhang et al., 2018)

**Figure 6: Figure Supplement 1.**
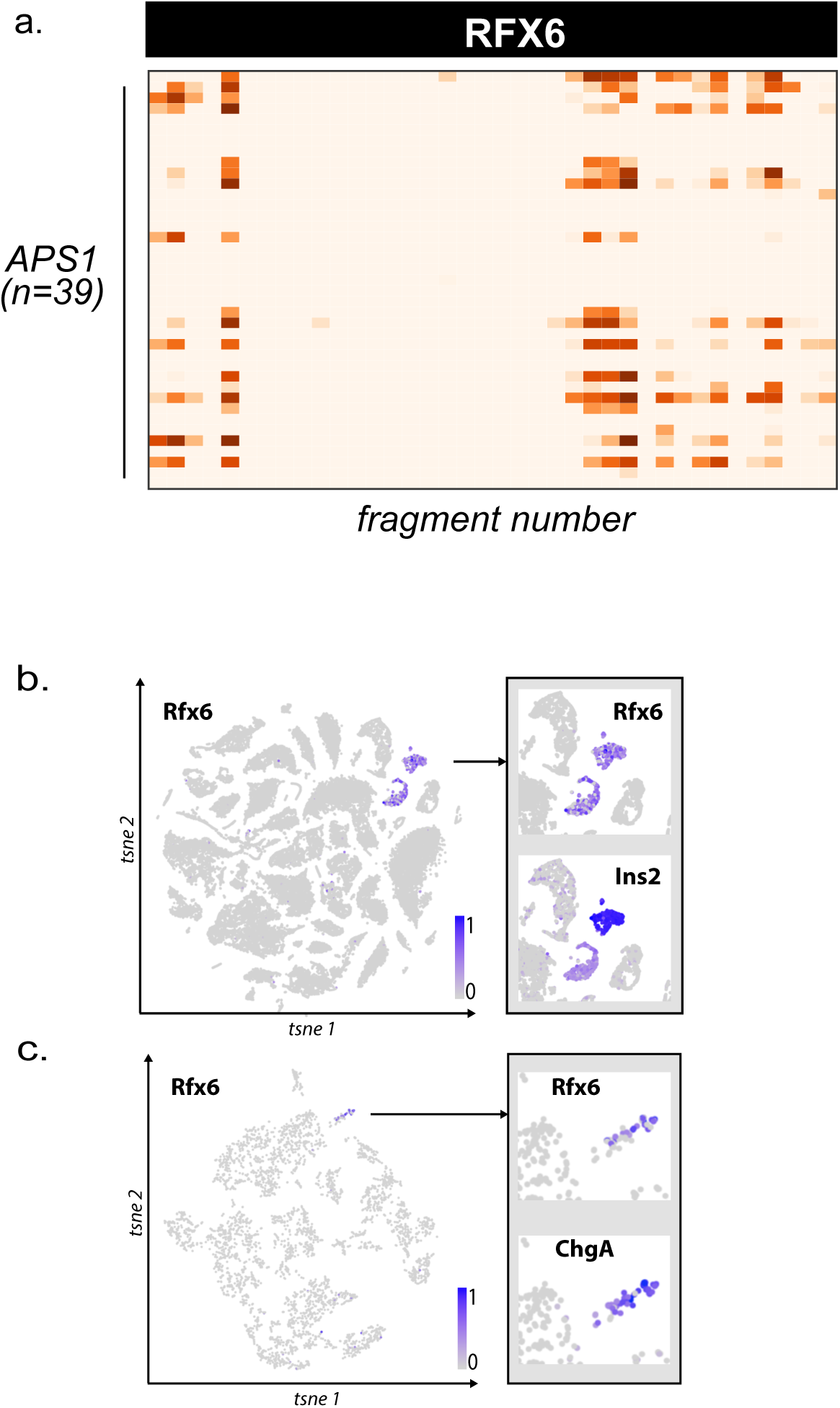
**A.** PhIP-Seq enables 49 amino acid resolution of antibody signal from novel autoantigen RFX6. PhIP-Seq signal (fold-change of each peptide as compared to signal from mock-IP, log10-scaled) for fragments 1-38 for RFX6 from APS1 sera (n=39). **B.** Single cell RNA expression of Rfx6. Left: normalized RNA expression of Rfx6 in single cells from 20 different organs. Right inset: Rfx6 shares an expression pattern with pancreatic beta-cell marker Ins2 (Schaum et al., 2018). **C.** Single cell RNA expression of Rfx6. Left: normalized RNA expression of Rfx6 in single cells from the intestine. Right inset: Rfx6 shares an expression pattern with intestinal enteroendocrine cell marker ChgA (Schaum et al., 2018).

**Figure 6: Figure Supplement 2.**
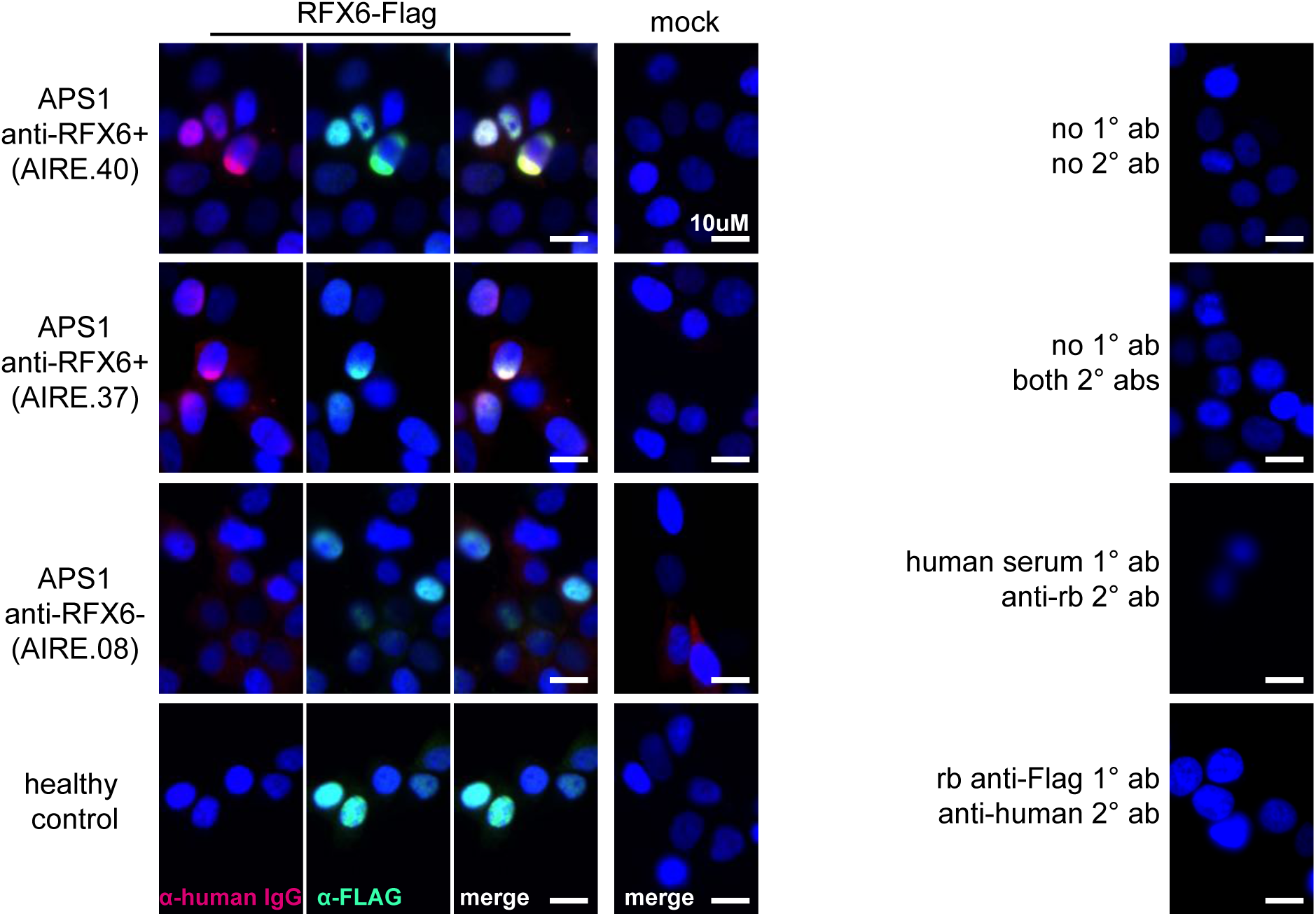
Anti-RFX6+ sera (top two panels), but not anti-RFX6-serum or non-APS1 control serum (bottom two panels), co-stain HEK293T cells transfected with an RFX6-expressing plasmid. None of the sera tested stain 293T cells transfected with empty vector (‘mock’). No cross-reactivity of secondary antibodies was observed (right panel).

**Figure 6: Figure Supplement 3.**
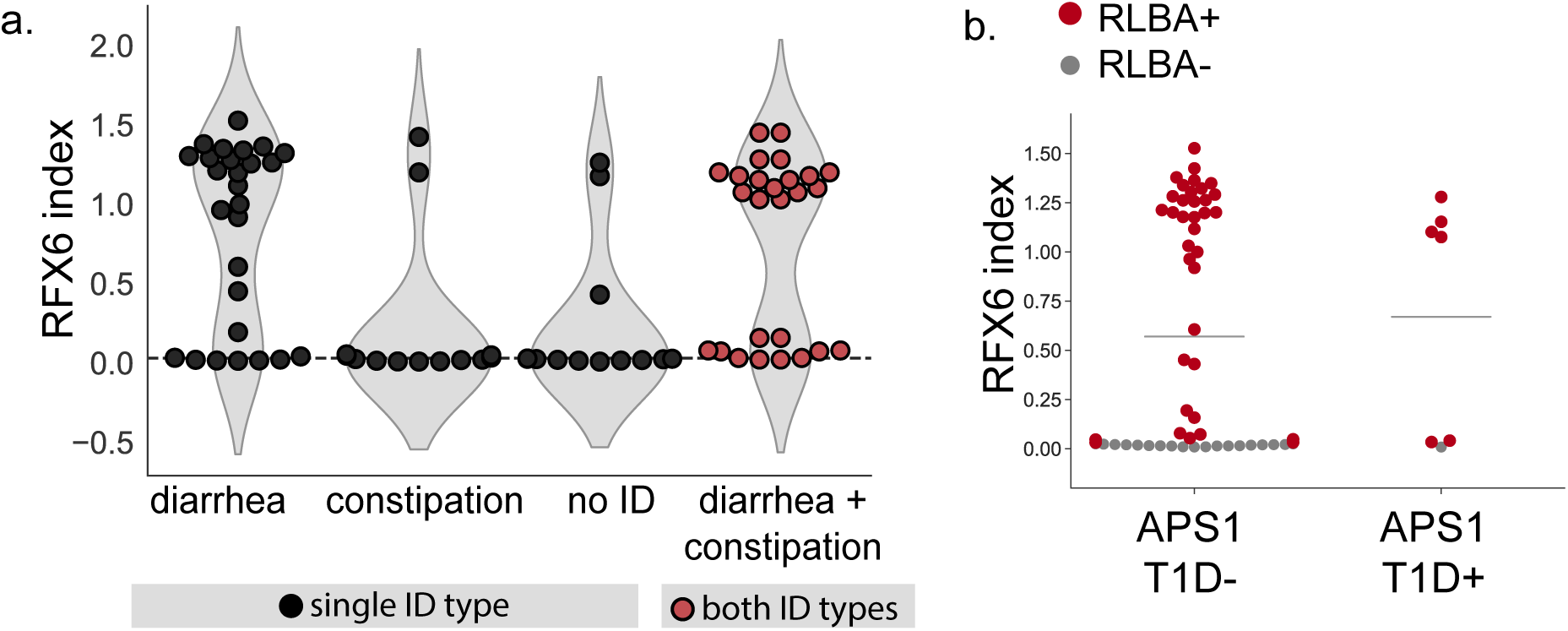
**A. APS1** patients with the diarrheal subtype, as well as those with both subtypes of ID (red), have increased anti-RFX6 antibody signal by RLBA as compared to those with constipation-type ID or no ID. B. 6/7 (6 diagnosed prior to serum draw, 1 diagnosed post serum draw) APS1 patients with type 1 diabetes have positive anti-RFX6 signal by RLBA.

## REFERENCES

Aaltonen, J., Björses, P., Perheentupa, J., Horelli–Kuitunen, N., Palotie, A., Peltonen, L., Lee, Y., Francis, F., Henning, S., Thiel, C., Leharach, H., & Yaspo, M. (1997). An autoimmune disease, APECED, caused by mutations in a novel gene featuring two PHD-type zinc-finger domains. Nature Genetics, 17(4), 399–403. https://doi.org/10.1038/ng1297-399

Ackermann, K., Bux, R., Rüb, U., Korf, H.-W., Kauert, G., & Stehle, J. H. (2006). Characterization of Human Melatonin Synthesis Using Autoptic Pineal Tissue. Endocrinology, 147(7), 3235–3242. https://doi.org/10.1210/en.2006-0043

Adriaenssens, A. E., Biggs, E. K., Darwish, T., Tadross, J., Sukthankar, T., Girish, M., Polex-Wolf, J., Lam, B. Y., Zvetkova, I., Pan, W., Chiarugi, D., Yeo, G., Blouet, C., Gribble, F. M., & Reimann, F. (2019). Glucose-Dependent Insulinotropic Polypeptide Receptor-Expressing Cells in the Hypothalamus Regulate Food Intake. Cell Metabolism. https://doi.org/10.1016/j.cmet.2019.07.013

Ahonen, P., Myllärniemi, S., Sipilä, I., & Perheentupa, J. (1990). Clinical Variation of Autoimmune Polyendocrinopathy–Candidiasis–Ectodermal Dystrophy (APECED) in a Series of 68 Patients. The New England Journal of Medicine, 322(26), 1829–1836. https://doi.org/10.1056/nejm199006283222601

Akoury, E., Zhang, L., Ao, A., & Slim, R. (2015). NLRP7 and KHDC3L, the two maternal-effect proteins responsible for recurrent hydatidiform moles, co-localize to the oocyte cytoskeleton. Human Reproduction (Oxford, England), 30(1), 159 169. https://doi.org/10.1093/humrep/deu291

Alanentalo, T., Chatonnet, F., Karlen, M., Sulniute, R., Ericson, J., Andersson, E., & Ahlgren, U. (2006). Cloning and analysis of Nkx6.3 during CNS and gastrointestinal development. Gene Expression Patterns, 6(2), 162–170. https://doi.org/10.1016/j.modgep.2005.06.012

Alimohammadi, M., Björklund, P., Hallgren, A., Pöntynen, N., Szinnai, G., Shikama, N., Keller, M. P., Ekwall, O., Kinkel, S. A., Husebye, E. S., Gustafsson, J., Rorsman, F., Peltonen, L., Betterle, C., Perheentupa, J., Akerström, G., Westin, G., Scott, H. S., Holländer, G. A., & Kämpe, O. (2008). Autoimmune polyendocrine syndrome type 1 and NALP5, a parathyroid autoantigen. The New England Journal of Medicine, 358(10), 1018 1028. https://doi.org/10.1056/nejmoa0706487

Alimohammadi, M., Dubois, N., Sköldberg, F., Hallgren, Å., Tardivel, I., Hedstrand, H., Haavik, J., Husebye, E. S., Gustafsson, J., Rorsman, F., Meloni, A., Janson, C., Vialettes, B., Kajosaari, M., Egner, W., Sargur, R., Pontén, F., Amoura, Z., Grimfeld, A., … Carel, J.-C. (2009). Pulmonary autoimmunity as a feature of autoimmune polyendocrine syndrome type 1 and identification of KCNRG as a bronchial autoantigen. Proceedings of the National Academy of Sciences, 106(11), 4396–4401. https://doi.org/10.1073/pnas.0809986106

Anderson. (2002). Projection of an Immunological Self Shadow Within the Thymus by the Aire Protein. Science, 298(5597), 1395 1401. https://doi.org/10.1126/science.1075958

Baekkeskov, S., Aanstoot, H., Christgau, S., Reetz, A., limena, Cascalho, M., Folli, F., Richter-Olesen, H., Camilli, D. P., & Camilli, P. (1990). Identification of the 64K autoantigen in insulin-dependent diabetes as the GABA-synthesizing enzyme glutamic acid decarboxylase. Nature, 347(6289), 151 156. https://doi.org/10.1038/347151a0

Bebbere, D., Masala, L., Albertini, D., & Ledda, S. (2016). The subcortical maternal complex: multiple functions for one biological structure? Journal of Assisted Reproduction and Genetics, 33(11), 1 8. https://doi.org/10.1007/s10815-016-0788-z

Berger, M., Scheel, D. W., Macias, H., Miyatsuka, T., Kim, H., Hoang, P., Ku, G. M., Honig, G., Liou, A., Tang, Y., Regard, J. B., Sharifnia, P., Yu, L., Wang, J., Coughlin, S. R., Conklin, B. R., Deneris, E. S., Tecott, L. H., & German, M. S. (2015). Gαi/o-coupled receptor signaling restricts pancreatic β-cell expansion. Proceedings of the National Academy of Sciences, 112(9), 2888–2893. https://doi.org/10.1073/pnas.1319378112

Berson, S. A., Yalow, R. S., Bauman, A., Rothschild, M. A., & Newerly, K. (1956). INSULIN-I131 METABOLISM IN HUMAN SUBJECTS: DEMONSTRATION OF INSULIN BINDING GLOBULIN IN THE CIRCULATION OF INSULIN TREATED SUBJECTS 1. Journal of Clinical Investigation, 35(2), 170–190. https://doi.org/10.1172/jci103262

Betterle, C., Pra, C., Mantero, F., & Zanchetta, R. (2002). Autoimmune Adrenal Insufficiency and Autoimmune Polyendocrine Syndromes: Autoantibodies, Autoantigens, and Their Applicability in Diagnosis and Disease Prediction. Endocrine Reviews, 23(3), 327–364. https://doi.org/10.1210/edrv.23.3.0466

Björk, E., Velloso, L. A., Kämpe, O., & Karlsson, A. F. (1994). GAD Autoantibodies in IDDM, Stiff-Man Syndrome, and Autoimmune Polyendocrine Syndrome Type I Recognize Different Epitopes. Diabetes, 43(1), 161–165. https://doi.org/10.2337/diab.43.1.161

Brozzetti, A., Alimohammadi, M., Morelli, S., Minarelli, V., Hallgren, A., Giordano, R., Bellis, A., Perniola, R., Kämpe, O., & Falorni, A. (2015). Autoantibody Response Against NALP5/MATER in Primary Ovarian Insufficiency and in Autoimmune Addison’s Disease. The Journal of Clinical Endocrinology & Metabolism, 100(5), 1941 1948. https://doi.org/10.1210/jc.2014-3571

Bruserud, Ø., Oftedal, B. E., Landegren, N., Erichsen, M. M., Bratland, E., Lima, K., Jørgensen, A. P., Myhre, A. G., Svartberg, J., Fougner, K. J., Bakke, Åsne, Nedrebø, B. G., Mella, B., Breivik, L., Viken, M. K., Knappskog, P. M., Marthinussen, M. C., Løvås, K., Kämpe, O., … Husebye, E. S. (2016). A Longitudinal Follow-up of Autoimmune Polyendocrine Syndrome Type 1. The Journal of Clinical Endocrinology & Metabolism, 101(8), 2975–2983. https://doi.org/10.1210/jc.2016-1821

Cheng, M., & Anderson, M. S. (2018). Thymic tolerance as a key brake on autoimmunity. Nature Immunology, 19(7), 659–664. https://doi.org/10.1038/s41590-018-0128-9

Choi, H., Kim, T.-H., Yun, C.-Y., Kim, J.-W., & Cho, E.-S. (2016). Testicular acid phosphatase induces odontoblast differentiation and mineralization. Cell and Tissue Research, 364(1), 95– 103. https://doi.org/10.1007/s00441-015-2310-9

Clemente, M., Obermayer-Straub, P., Meloni, A., Strassburg, C. P., Arangino, V., Tukey, R. H., Virgiliis, S., & Manns, M. P. (1997). Cytochrome P450 1A2 Is a Hepatic Autoantigen in Autoimmune Polyglandular Syndrome Type 1 1. The Journal of Clinical Endocrinology & Metabolism, 82(5), 1353–1361. https://doi.org/10.1210/jcem.82.5.3913

Conteduca, G., Indiveri, F., Filaci, G., & Negrini, S. (2018). Beyond APECED: An update on the role of the AutoImmune regulator gene (AIRE) in physiology and disease. Autoimmunity Reviews, 17(Immunity 44 2016), 1 17. https://doi.org/10.1016/j.autrev.2017.10.017

DeVoss, J., Shum, A., Johannes, K., Lu, W., Krawisz, A., Wang, P., Yang, T., LeClair, N., Austin, C., Strauss, E., & Anderson. (2008). Effector Mechanisms of the Autoimmune Syndrome in the Murine Model of Autoimmune Polyglandular Syndrome Type 1. The Journal of Immunology, 181(6), 4072 4079. https://doi.org/10.4049/jimmunol.181.6.4072

Du, Y. T., Rayner, C. K., Jones, K. L., Talley, N. J., & Horowitz, M. (2018). Gastrointestinal Symptoms in Diabetes: Prevalence, Assessment, Pathogenesis, and Management. Diabetes Care, 41(3), 627–637. https://doi.org/10.2337/dc17-1536

Ekwall, O., Hedstrand, H., Grimelius, L., Haavik, J., Perheentupa, J., Gustafsson, J., Husebye, E., Kämpe, O., & Rorsman, F. (1998). Identification of tryptophan hydroxylase as an intestinal autoantigen. The Lancet, 352(9124), 279–283. https://doi.org/10.1016/s0140-6736(97)11050-9

Ferré, E. M., Break, T. J., Burbelo, P. D., Allgäuer, M., Kleiner, D. E., Jin, D., Xu, Z., Folio, L. R., Mollura, D. J., Swamydas, M., Gu, W., Hunsberger, S., Lee, C.-C. R., Bondici, A., Hoffman, K. W., Lim, J. K., Dobbs, K., Niemela, J. E., Fleisher, T. A., … Lionakis, M. S. (2019). Lymphocyte-driven regional immunopathology in pneumonitis caused by impaired central immune tolerance. Science Translational Medicine, 11(495), eaav5597. https://doi.org/10.1126/scitranslmed.aav5597

Ferré, E. M., Rose, S. R., Rosenzweig, S. D., Burbelo, P. D., Romito, K. R., Niemela, J. E., Rosen, L. B., Break, T. J., Gu, W., Hunsberger, S., Browne, S. K., Hsu, A. P., Rampertaap, S., Swamydas, M., Collar, A. L., Kong, H. H., Lee, C.-C., Chascsa, D., Simcox, T., … Lionakis, M. S. (2016). Redefined clinical features and diagnostic criteria in autoimmune polyendocrinopathy-candidiasis-ectodermal dystrophy. JCI Insight, 1(13), 1343 19. https://doi.org/10.1172/jci.insight.88782

Fishman, D., Kisand, K., Hertel, C., Rothe, M., Remm, A., Pihlap, M., Adler, P., Vilo, J., Peet, A., Meloni, A., Podkrajsek, K., Battelino, T., Bruserud, Ø., Wolff, A. S., Husebye, E. S., Kluger, N., Krohn, K., Ranki, A., Peterson, H., … Peterson, P. (2017). Autoantibody Repertoire in APECED Patients Targets Two Distinct Subgroups of Proteins. Frontiers in Immunology, 8, 681 15. https://doi.org/10.3389/fimmu.2017.00976

Gavanescu, I., Benoist, C., & Mathis, D. (2008). B cells are required for Aire-deficient mice to develop multi-organ autoinflammation: A therapeutic approach for APECED patients. Proceedings of the National Academy of Sciences, 105(35), 13009–13014. https://doi.org/10.1073/pnas.0806874105

Gehart, H., van Es, J. H., Hamer, K., Beumer, J., Kretzschmar, K., Dekkers, J. F., Rios, A., & Clevers, H. (2019). Identification of Enteroendocrine Regulators by Real-Time Single-Cell Differentiation Mapping. Cell, 176(Dev. Dyn. 194 1992), 1158–1173.e16. https://doi.org/10.1016/j.cell.2018.12.029

Goldspink, D. A., Reimann, F., & Gribble, F. M. (2018). Models and Tools for Studying Enteroendocrine Cells. Endocrinology, 159(12), 3874–3884. https://doi.org/10.1210/en.2018-00672

Hedstrand, H., Ekwall, O., Olsson, M. J., Landgren, E., Kemp, H. E., Weetman, A. P., Perheentupa, J., Husebye, E., Gustafsson, J., Betterle, C., Kämpe, O., & Rorsman, F. (2001). The Transcription Factors SOX9 and SOX10 Are Vitiligo Autoantigens in Autoimmune Polyendocrine Syndrome Type I. Journal of Biological Chemistry, 276(38), 35390 35395. https://doi.org/10.1074/jbc.m102391200

Högenauer, C., Meyer, R. L., Netto, G. J., Bell, D., Little, K. H., Ferries, L., Ana, C. A., Porter, J. L., & Fordtran, J. S. (2001). Malabsorption Due to Cholecystokinin Deficiency in a Patient with Autoimmune Polyglandular Syndrome Type I. The New England Journal of Medicine, 344(4), 270–274. https://doi.org/10.1056/nejm200101253440405

Holland, A. M., Hale, M. A., Kagami, H., Hammer, R. E., & MacDonald, R. J. (2002). Experimental control of pancreatic development and maintenance. Proceedings of the National Academy of Sciences, 99(19), 12236–12241. https://doi.org/10.1073/pnas.192255099

Huhtaniemi, I., Hovatta, O., Marca, A., Livera, G., Monniaux, D., Persani, L., Heddar, A., Jarzabek, K., Laisk-Podar, T., Salumets, A., Tapanainen, J. S., Veitia, R. A., Visser, J. A., Wieacker, P., Wolczynski, S., & Misrahi, M. (2018). Advances in the Molecular Pathophysiology, Genetics, and Treatment of Primary Ovarian Insufficiency. Trends in Endocrinology & Metabolism, 29(6), 400 419. https://doi.org/10.1016/j.tem.2018.03.010

Husebye, E. S., Anderson, M. S., & Kämpe, O. (2018). Autoimmune Polyendocrine Syndromes. The New England Journal of Medicine, 378(12), 1132–1141. https://doi.org/10.1056/nejmra1713301

Husebye, E. S., Gebre-Medhin, G., Tuomi, T., Perheentupa, J., Landin-Olsson, M., Gustafsson, J., Rorsman, F., & Kämpe, O. (1997). Autoantibodies against Aromatic l -Amino Acid Decarboxylase in Autoimmune Polyendocrine Syndrome Type I 1. The Journal of Clinical Endocrinology & Metabolism, 82(1), 147–150. https://doi.org/10.1210/jcem.82.1.3647

Jasti, S., Warren, B. D., McGinnis, L. K., Kinsey, W. H., Petroff, B. K., & Petroff, M. G. (2012). The Autoimmune Regulator Prevents Premature Reproductive Senescence in Female Mice1. Biology of Reproduction, 86(4), 163 9. https://doi.org/10.1095/biolreprod.111.097501

Jeong, J., Jiang, L., Albino, E., Marrero, J., Rho, H., Hu, J., Hu, S., Vera, C., Bayron-Poueymiroy, D., Rivera-Pacheco, Z., Ramos, L., Torres-Castro, C., Qian, J., Bonaventura, J., Boeke, J. D., Yap, W. Y., Pino, I., Eichinger, D. J., Zhu, H., & Blackshaw, S. (2012). Rapid Identification of Monospecific Monoclonal Antibodies Using a Human Proteome Microarray. Molecular & Cellular Proteomics, 11(6), O111.016253. https://doi.org/10.1074/mcp.o111.016253

Kluger, N., Jokinen, M., Lintulahti, A., Krohn, K., & Ranki, A. (2015). Gastrointestinal immunity against tryptophan hydroxylase-1, aromatic L-amino-acid decarboxylase, AIE-75, villin and Paneth cells in APECED. Clinical Immunology, 158(2), 212–220. https://doi.org/10.1016/j.clim.2015.03.012

Kuroda, N., Mitani, T., Takeda, N., Ishimaru, N., Arakaki, R., Hayashi, Y., Bando, Y., Izumi, K., Takahashi, T., Nomura, T., Sakaguchi, S., Ueno, T., Takahama, Y., Uchida, D., Sun, S., Kajiura, F., Mouri, Y., Han, H., Matsushima, A., … Matsumoto, M. (2005). Development of Autoimmunity against Transcriptionally Unrepressed Target Antigen in the Thymus of Aire-Deficient Mice. The Journal of Immunology, 174(4), 1862–1870. https://doi.org/10.4049/jimmunol.174.4.1862

Landegren, N., Sharon, D., Freyhult, E., Hallgren, A., Eriksson, D., Edqvist, P.-H., Bensing, S., Wahlberg, J., Nelson, L. M., Gustafsson, J., Husebye, E. S., Anderson, M. S., Snyder, M., & Kämpe, O. (2016). Proteome-wide survey of the autoimmune target repertoire in autoimmune polyendocrine syndrome type 1. Nature Publishing Group, 6(1), 1 11. https://doi.org/10.1038/srep20104

Landegren, N., Sharon, D., Shum, A. K., Khan, I. S., Fasano, K. J., Hallgren, Å., Kampf, C., Freyhult, E., Ardesjö-Lundgren, B., Alimohammadi, M., Rathsman, S., Ludvigsson, J. F., Lundh, D., Motrich, R., Rivero, V., Fong, L., Giwercman, A., Gustafsson, J., Perheentupa, J., … Kämpe, O. (2015). Transglutaminase 4 as a prostate autoantigen in male subfertility. Science Translational Medicine, 7(292), 292ra101–292ra101. https://doi.org/10.1126/scitranslmed.aaa9186

Lanzavecchia, A. (1985). Antigen-specific interaction between T and B cells. Nature, 314(6011), 537–539. https://doi.org/10.1038/314537a0

Larman, B. H., Zhao, Z., Laserson, U., Li, M. Z., Ciccia, A., Gakidis, A. M., Church, G. M., Kesari, S., LeProust, E. M., Solimini, N. L., & Elledge, S. J. (2011). Autoantigen discovery with a synthetic human peptidome. Nature Biotechnology, 29(6), 535 541. https://doi.org/10.1038/nbt.1856

Leonard, J. D., Gilmore, D. C., Dileepan, T., Nawrocka, W. I., Chao, J. L., Schoenbach, M. H., Jenkins, M. K., Adams, E. J., & Savage, P. A. (2017). Identification of Natural Regulatory T Cell Epitopes Reveals Convergence on a Dominant Autoantigen. Immunity, 47(1), 107–117.e8. https://doi.org/10.1016/j.immuni.2017.06.015

Li, L., Baibakov, B., & Dean, J. (2008). A Subcortical Maternal Complex Essential for Preimplantation Mouse Embryogenesis. Developmental Cell, 15(3), 416 425. https://doi.org/10.1016/j.devcel.2008.07.010

Liu, C., Li, M., Li, T., Zhao, H., Huang, J., Wang, Y., Gao, Q., Yu, Y., & Shi, Q. (2016). ECAT1 is essential for human oocyte maturation and pre-implantation development of the resulting embryos. Scientific Reports, 6(1), 38192. https://doi.org/10.1038/srep38192

Maclaren, N., Chen, Q., Kukreja, A., Marker, J., Zhang, C., & Sun, Z. (2001). Autoimmune hypogonadism as part of an autoimmune polyglandular syndrome. Journal of the Society for Gynecologic Investigation, 8(1), S52–S54. https://doi.org/10.1016/s1071-5576(00)00109-x

Malchow, S., Leventhal, D. S., Lee, V., Nishi, S., Socci, N. D., & Savage, P. A. (2016). Aire Enforces Immune Tolerance by Directing Autoreactive T Cells into the Regulatory T Cell Lineage. Immunity, 44(5), 1102 1113. https://doi.org/10.1016/j.immuni.2016.02.009

Mandel-Brehm, C., Dubey, D., Kryzer, T. J., O’Donovan, B. D., Tran, B., Vazquez, S. E., Sample, H. A., Zorn, K. C., Khan, L. M., Bledsoe, I. O., McKeon, A., Pleasure, S. J., Lennon, V. A., DeRisi, J. L., Wilson, M. R., & Pittock, S. J. (2019). Kelch-like Protein 11 Antibodies in Seminoma-Associated Paraneoplastic Encephalitis. New England Journal of Medicine, 381(1), 47–54. https://doi.org/10.1056/nejmoa1816721

Meager, A., Visvalingam, K., Peterson, P., Möll, K., Murumägi, A., Krohn, K., Eskelin, P., Perheentupa, J., Husebye, E., Kadota, Y., & Willcox, N. (2006). Anti-Interferon Autoantibodies in Autoimmune Polyendocrinopathy Syndrome Type 1. PLoS Medicine, 3(7), e289. https://doi.org/10.1371/journal.pmed.0030289

Meyer, S., Woodward, M., Hertel, C., Vlaicu, P., Haque, Y., Kärner, J., Macagno, A., Onuoha, S. C., Fishman, D., Peterson, H., Metsküla, K., Uibo, R., Jäntti, K., Hokynar, K., Wolff, A. S., patient collaborative, A., Meloni, A., Kluger, N., Husebye, E. S., … Hayday, A. (2016). AIRE-Deficient Patients Harbor Unique High-Affinity Disease-Ameliorating Autoantibodies. Cell, 166(3), 582 595. https://doi.org/10.1016/j.cell.2016.06.024

Mitchell, J., Punthakee, Z., Lo, B., Bernard, C., Chong, K., Newman, C., Cartier, L., Desilets, V., Cutz, E., Hansen, I., Riley, P., & Polychronakos, C. (2004). Neonatal diabetes, with hypoplastic pancreas, intestinal atresia and gall bladder hypoplasia: search for the aetiology of a new autosomal recessive syndrome. Diabetologia, 47(12), 2160 2167. https://doi.org/10.1007/s00125-004-1576-3

Moody, A. J., Thim, L., & Valverde, I. (1984). The isolation and sequencing of human gastric inhibitory peptide (GIP). FEBS Letters, 172(2), 142–148. https://doi.org/10.1016/0014-5793(84)81114-x

Nagamine, K., Peterson, P., Scott, H. S., Kudoh, J., Minoshima, S., Heino, M., Krohn, K. J., Lalioti, M. D., Mullis, P. E., Antonarakis, S. E., Kawasaki, K., Asakawa, S., Ito, F., & Shimizu, N. (1997). Positional cloning of the APECED gene. Nature Genetics, 17(4), 393–398. https://doi.org/10.1038/ng1297-393

Nelson, L. M. (2009). Clinical practice. Primary ovarian insufficiency. The New England Journal of Medicine, 360(6), 606 614. https://doi.org/10.1056/nejmcp0808697

Obermayer-Straub, P., Strassburg, C., & Manns, M. (2000). Target Proteins in Human Autoimmunity: Cytochromes P450 and Udp-Glycoronosyltransferases. Canadian Journal of Gastroenterology and Hepatology, 14(5), 429–439. https://doi.org/10.1155/2000/910107

O’Connor, D. T., Burton, D., & Deftos, L. J. (1983). Chromogranin A: Immunohistology reveals its universal occurrence in normal polypeptide hormone producing endocrine glands. Life Sciences, 33(17), 1657–1663. https://doi.org/10.1016/0024-3205(83)90721-x

O’Donovan, B., Mandel - Brehm, C., E. Vazquez, S., Liu, J., Paren, A. V., Anderson, M. S., Kassimatis, T., Zekeridou, A., Hauser, S. L., Pittock, S. J., Chow, E., Wilson, M. R., & DeRisi, J. L. (2018). Exploration of Anti - Yo and Anti - Hu paraneoplastic neurological disorders by PhIP - Seq reveals a highly restricted pattern of antibody epitopes. Biorxiv.Org. https://doi.org/10.1101/502187

Oftedal, B. E., Hellesen, A., Erichsen, M. M., Bratland, E., Vardi, A., Perheentupa, J., Kemp, H. E., Fiskerstrand, T., Viken, M. K., Weetman, A. P., Fleishman, S. J., Banka, S., Newman, W. G., Sewell, W. A. C., Sozaeva, L. S., Zayats, T., Haugarvoll, K., Orlova, E. M., Haavik, J., … Husebye, E. S. (2015). Dominant Mutations in the Autoimmune Regulator AIRE Are Associated with Common Organ-Specific Autoimmune Diseases. Immunity, 42(6), 1185–1196. https://doi.org/10.1016/j.immuni.2015.04.021

Ohsie, S., Gerney, G., Gui, D., Kahana, D., Martín, M. G., & Cortina, G. (2009). A paucity of colonic enteroendocrine and/or enterochromaffin cells characterizes a subset of patients with chronic unexplained diarrhea/malabsorption. Human Pathology, 40(7), 1006–1014. https://doi.org/10.1016/j.humpath.2008.12.016

Oliva-Hemker, M., Berkenblit, G., Anhalt, G., & Yardley, J. (2006). Pernicious Anemia and Widespread Absence of Gastrointestinal Endocrine Cells in a Patient with Autoimmune Polyglandular Syndrome Type I and Malabsorption. The Journal of Clinical Endocrinology & Metabolism, 91(8), 2833–2838. https://doi.org/10.1210/jc.2005-2506

Otsuka, N., Tong, Z.-B., Vanevski, K., Tu, W., Cheng, M. H., & Nelson, L. M. (2011). Autoimmune Oophoritis with Multiple Molecular Targets Mitigated by Transgenic Expression of Mater. Endocrinology, 152(6), 2465 2473. https://doi.org/10.1210/en.2011-0022

Patel, K. A., Kettunen, J., Laakso, M., Stančáková, A., Laver, T. W., Colclough, K., Johnson, M. B., Abramowicz, M., Groop, L., Miettinen, P. J., Shepherd, M. H., Flanagan, S. E., Ellard, S., Inagaki, N., Hattersley, A. T., Tuomi, T., Cnop, M., & Weedon, M. N. (2017). Heterozygous RFX6 protein truncating variants are associated with MODY with reduced penetrance. Nature Communications, 8(1), 1 8. https://doi.org/10.1038/s41467-017-00895-9

Pederson, R. A., & McIntosh, C. H. (2016). Discovery of gastric inhibitory polypeptide and its subsequent fate: Personal reflections. Journal of Diabetes Investigation, 7(S1), 4–7. https://doi.org/10.1111/jdi.12480

Piccand, J., Vagne, C., Blot, F., Meunier, A., Beucher, A., Strasser, P., Lund, M. L., Ghimire, S., Nivlet, L., Lapp, C., Petersen, N., Engelstoft, M. S., Thibault-Carpentier, C., Keime, C., Correa, S., Schreiber, V., Molina, N., Schwartz, T. W., Arcangelis, A., & Gradwohl, G. (2019). Rfx6 promotes the differentiation of peptide-secreting enteroendocrine cells while repressing genetic programs controlling serotonin production. Molecular Metabolism, 29, 24–39. https://doi.org/10.1016/j.molmet.2019.08.007

Pöntynen, N., Miettinen, A., Arstila, P. T., Kämpe, O., Alimohammadi, M., Vaarala, O., Peltonen, L., & Ulmanen, I. (2006). Aire deficient mice do not develop the same profile of tissue-specific autoantibodies as APECED patients. Journal of Autoimmunity, 27(2), 96–104. https://doi.org/10.1016/j.jaut.2006.06.001

Popler, J., Alimohammadi, M., Kämpe, O., Dalin, F., Dishop, M. K., Barker, J. M., Moriarty-Kelsey, M., Soep, J. B., & Deterding, R. R. (2012). Autoimmune polyendocrine syndrome type 1: Utility of KCNRG autoantibodies as a marker of active pulmonary disease and successful treatment with rituximab. Pediatric Pulmonology, 47(1), 84–87. https://doi.org/10.1002/ppul.21520

Posovszky, C., Lahr, G., von Schnurbein, J., Buderus, S., Findeisen, A., Schröder, C., Schütz, C., Schulz, A., batin, K., Wabitsch, M., & Barth, T. (2012). Loss of Enteroendocrine Cells in Autoimmune-Polyendocrine-Candidiasis-Ectodermal-Dystrophy (APECED) Syndrome with Gastrointestinal Dysfunction. The Journal of Clinical Endocrinology & Metabolism, 97(2), E292 E300. https://doi.org/10.1210/jc.2011-2044

Puel, A., Döffinger, R., Natividad, A., Chrabieh, M., Barcenas-Morales, G., Picard, C., Cobat, A., Ouachée-Chardin, M., Toulon, A., Bustamante, J., Al-Muhsen, S., Al-Owain, M., Arkwright, P. D., Costigan, C., McConnell, V., Cant, A. J., Abinun, M., Polak, M., Bougnères, P.-F., … Casanova, J.-L. (2010). Autoantibodies against IL-17A, IL-17F, and IL-22 in patients with chronic mucocutaneous candidiasis and autoimmune polyendocrine syndrome type I. The Journal of Experimental Medicine, 207(2), 291–297. https://doi.org/10.1084/jem.20091983

Rath, M. F., Coon, S. L., Amaral, F. G., Weller, J. L., Møller, M., & Klein, D. C. (2016). Melatonin Synthesis: Acetylserotonin O-Methyltransferase (ASMT) Is Strongly Expressed in a Subpopulation of Pinealocytes in the Male Rat Pineal Gland. Endocrinology, 157(5), 2028–2040. https://doi.org/10.1210/en.2015-1888

Reddy, R., Akoury, E., Nguyen, N., Abdul-Rahman, O. A., Dery, C., Gupta, N., Daley, W. P., Ao, A., Landolsi, H., Fisher, R., Touitou, I., & Slim, R. (2012). Report of four new patients with protein-truncating mutations in C6orf221/KHDC3L and colocalization with NLRP7. European Journal of Human Genetics, 21(9), 957 964. https://doi.org/10.1038/ejhg.2012.274

Rezaei, M., Nguyen, N., Foroughinia, L., Dash, P., Ahmadpour, F., Verma, I., Slim, R., & Fardaei, M. (2016). Two novel mutations in the KHDC3L gene in Asian patients with recurrent hydatidiform mole. Human Genome Variation, 3(1), 16027 5. https://doi.org/10.1038/hgv.2016.27

Rosen, A., & Casciola-Rosen, L. (2014). Autoantigens as Partners in Initiation and Propagation of Autoimmune Rheumatic Diseases. Annual Review of Immunology, 34(1), 1–26. https://doi.org/10.1146/annurev-immunol-032414-112205

Sansom, S. N., Shikama-Dorn, N., Zhanybekova, S., Nusspaumer, G., Macaulay, I. C., Deadman, M. E., Heger, A., Ponting, C. P., & Holländer, G. A. (2014). Population and single-cell genomics reveal the Aire dependency, relief from Polycomb silencing, and distribution of self-antigen expression in thymic epithelia. Genome Research, 24(12), 1918–1931. https://doi.org/10.1101/gr.171645.113

Schaum, N., Karkanias, J., Neff, N. F., May, A. P., Quake, S. R., Wyss-Coray, T., Darmanis, S., Batson, J., Botvinnik, O., Chen, M. B., Chen, S., Green, F., Jones, R. C., Maynard, A., Penland, L., Pisco, A., Sit, R. V., anley, G., Webber, J. T., … Wyss-Coray, T. (2018). Single-cell transcriptomics of 20 mouse organs creates a Tabula Muris. Nature, 562(7727), 1 25. https://doi.org/10.1038/s41586-018-0590-4

Seymen, F., Kim, Y., Lee, Y., Kang, J., Kim, T.-H., Choi, H., Koruyucu, M., Kasimoglu, Y., Tuna, E., Gencay, K., Shin, T., Hyun, H.-K., Kim, Y.-J., Lee, S.-H., Lee, Z., Zhang, H., Hu, J., Simmer, J. P., Cho, E.-S., & Kim, J.-W. (2016). Recessive Mutations in ACPT, Encoding Testicular Acid Phosphatase, Cause Hypoplastic Amelogenesis Imperfecta. The American Journal of Human Genetics, 99(5), 1199–1205. https://doi.org/10.1016/j.ajhg.2016.09.018

Sharon, D., & Snyder, M. (2014). Serum Profiling Using Protein Microarrays to Identify Disease Related Antigens. Methods in Molecular Biology (Clifton, N.J.), 1176, 169–178. https://doi.org/10.1007/978-1-4939-0992-6_14

Shum, A. K., Alimohammadi, M., Tan, C. L., Cheng, M. H., Metzger, T. C., Law, C. S., Lwin, W., Perheentupa, J., Bour-Jordan, H., Carel, J., Husebye, E. S., Luca, F., Janson, C., Sargur, R., Dubois, N., Kajosaari, M., Wolters, P. J., Chapman, H. A., Kämpe, O., & Anderson, M. S. (2013). BPIFB1 is a lung-specific autoantigen associated with interstitial lung disease. Science Translational Medicine, 5(206), 206ra139 206ra139. https://doi.org/10.1126/scitranslmed.3006998

Shum, A. K., DeVoss, J., Tan, C. L., Hou, Y., Johannes, K., O’Gorman, C. S., Jones, K. D., Sochett, E. B., Fong, L., & Anderson, M. S. (2009). Identification of an autoantigen demonstrates a link between interstitial lung disease and a defect in central tolerance. Science Translational Medicine, 1(9), 9ra20 9ra20. https://doi.org/10.1126/scitranslmed.3000284

Sifuentes-Dominguez, L. F., Li, H., Llano, E., Liu, Z., Singla, A., Patel, A. S., Kathania, M., Khoury, A., Norris, N., Rios, J. J., Starokadomskyy, P., Park, J. Y., Gopal, P., Liu, Q., Tan, S., Chan, L., Ross, T., Harrison, S., Venuprasad, K., … Burstein, E. (2019). SCGN deficiency results in colitis susceptibility. ELife, 8, e49910. https://doi.org/10.7554/elife.49910

Silva, C., Yamakami, L., Aikawa, N., aujo, D., Carvalho, J., & Bonfá, E. (2014). Autoimmune primary ovarian insufficiency. Autoimmunity Reviews, 13(4–5), 427 430. https://doi.org/10.1016/j.autrev.2014.01.003

Smith, C. E., Whitehouse, L. L., Poulter, J. A., Brookes, S. J., Day, P. F., Soldani, F., Kirkham, J., Inglehearn, C. F., & Mighell, A. J. (2017). Defects in the acid phosphatase ACPT cause recessive hypoplastic amelogenesis imperfecta. European Journal of Human Genetics, 25(8), 1015. https://doi.org/10.1038/ejhg.2017.79

Smith, S. B., Qu, H.-Q., Taleb, N., Kishimoto, N. Y., Scheel, D. W., Lu, Y., Patch, A.-M., Grabs, R., Wang, J., Lynn, F. C., Miyatsuka, T., Mitchell, J., Seerke, R., Désir, J., Eijnden, S., Abramowicz, M., Kacet, N., Weill, J., Renard, M.-È., … German, M. S. (2010). Rfx6 directs islet formation and insulin production in mice and humans. Nature, 463(7282), 775 780. https://doi.org/10.1038/nature08748

Sng, J., Ayoglu, B., Chen, J. W., Schickel, J.-N., Ferré, E. M., Glauzy, S., Romberg, N., Hoenig, M., Cunningham-Rundles, C., Utz, P. J., Lionakis, M. S., & Meffre, E. (2019). AIRE expression controls the peripheral selection of autoreactive B cells. Science Immunology, 4(34), eaav6778. https://doi.org/10.1126/sciimmunol.aav6778

Söderbergh, A., Myhre, A., Ekwall, O., Gebre-Medhin, G., Hedstrand, H., Landgren, E., Miettinen, A., Eskelin, P., Halonen, M., Tuomi, T., Gustafsson, J., Husebye, E. S., Perheentupa, J., Gylling, M., Manns, M. P., Rorsman, F., Kämpe, O., & Nilsson, T. (2004). Prevalence and Clinical Associations of 10 Defined Autoantibodies in Autoimmune Polyendocrine Syndrome Type I. The Journal of Clinical Endocrinology & Metabolism, 89(2), 557–562. https://doi.org/10.1210/jc.2003-030279

Stoffers, D. A., Zinkin, N. T., Stanojevic, V., Clarke, W. L., & Habener, J. F. (1997). Pancreatic agenesis attributable to a single nucleotide deletion in the human IPF1 gene coding sequence. Nature Genetics, 15(1), 106–110. https://doi.org/10.1038/ng0197-106

Taplin, C. E., & Barker, J. M. (2009). Autoantibodies in type 1 diabetes. Autoimmunity, 41(1), 11 18. https://doi.org/10.1080/08916930701619169

Uhlen, M., Fagerberg, L., Hallstrom, B., Lindskog, C., Oksvold, P., Mardinoglu, A., Sivertsson, A., Kampf, C., Sjostedt, E., Asplund, A., Olsson, I., Edlund, K., Lundberg, E., Navani, S., Szigyarto, C., Odeberg, J., Djureinovic, D., Takanen, J., Hober, S., … Ponten, F. (2015). Tissue-based map of the human proteome. Science, 347(6220), 1260419 1260419. https://doi.org/10.1126/science.1260419

Virant-Klun, I., Leicht, S., Hughes, C., & Krijgsveld, J. (2016). Identification of Maturation-Specific Proteins by Single-Cell Proteomics of Human Oocytes. Molecular & Cellular Proteomics : MCP, 15(8), 2616 2627. https://doi.org/10.1074/mcp.m115.056887

Wang, J., Cortina, G., Wu, V. S., Tran, R., Cho, J.-H., Tsai, M.-J., Bailey, T. J., Jamrich, M., Ament, M. E., Treem, W. R., Hill, I. D., Vargas, J. H., Gershman, G., Farmer, D. G., Reyen, L., & Martín, M. G. (2006). Mutant Neurogenin-3 in Congenital Malabsorptive Diarrhea. The New England Journal of Medicine, 355(3), 270–280. https://doi.org/10.1056/nejmoa054288

Wang, X., Song, D., Mykytenko, D., Kuang, Y., Lv, Q., Li, B., Chen, B., Mao, X., Xu, Y., Zukin, V., Mazur, P., Mu, J., Yan, Z., Zhou, Z., Li, Q., Liu, S., Jin, L., He, L., Sang, Q., … Wang, L. (2018). Novel mutations in genes encoding subcortical maternal complex proteins may cause human embryonic developmental arrest. Reproductive BioMedicine Online, 36(6), 698–704. https://doi.org/10.1016/j.rbmo.2018.03.009

Welt, C. K. (2008). Autoimmune Oophoritis in the Adolescent. Annals of the New York Academy of Sciences, 1135(1), 118–122. https://doi.org/10.1196/annals.1429.006

Winqvist, O., Gustafsson, J., Rorsman, F., Karlsson, F., & Kämpe, O. (1993). Two different cytochrome P450 enzymes are the adrenal antigens in autoimmune polyendocrine syndrome type I and Addison’s disease. Journal of Clinical Investigation, 92(5), 2377–2385. https://doi.org/10.1172/jci116843

Wolff, A., Sarkadi, A., Maródi, L., Kärner, J., Orlova, E., Oftedal, B., Kisand, K., Oláh, É., Meloni, A., Myhre, A., Husebye, E., Motaghedi, R., Perheentupa, J., Peterson, P., Willcox, N., & Meager, A. (2013). Anti-Cytokine Autoantibodies Preceding Onset of Autoimmune Polyendocrine Syndrome Type I Features in Early Childhood. Journal of Clinical Immunology, 33(8), 1341–1348. https://doi.org/10.1007/s10875-013-9938-6

Zhang, W., Chen, Z., Zhang, D., Zhao, B., Liu, L., Xie, Z., Yao, Y., & Zheng, P. (2019). KHDC3L mutation causes recurrent pregnancy loss by inducing genomic instability of human early embryonic cells. PLOS Biology, 17(10), e3000468. https://doi.org/10.1371/journal.pbio.3000468

Zhang, Y., Yan, Z., Qin, Q., Nisenblat, V., Chang, H.-M., Yu, Y., Wang, T., Lu, C., Yang, M., Yang, S., Yao, Y., Zhu, X., Xia, X., Dang, Y., Ren, Y., Yuan, P., Li, R., Liu, P., Guo, H., … Yan, L. (2018). Transcriptome Landscape of Human Folliculogenesis Reveals Oocyte and Granulosa Cell Interactions. Molecular Cell, 72(6), 1 19. https://doi.org/10.1016/j.molcel.2018.10.029

Zhu, H., Bilgin, M., Bangham, R., Hall, D., Casamayor, A., Bertone, P., Lan, N., Jansen, R., Bidlingmaier, S., Houfek, T., Mitchell, T., Miller, P., Dean, R. A., Gerstein, M., & Snyder, M. (2001). Global Analysis of Protein Activities Using Proteome Chips. Science, 293(5537), 2101–2105. https://doi.org/10.1126/science.1062191

Zhu, K., Yan, L., Zhang, X., Lu, X., Wang, T., Yan, J., Liu, X., Qiao, J., & Li, L. (2015). Identification of a human subcortical maternal complex. Molecular Human Reproduction, 21(4), 320 329. https://doi.org/10.1093/molehr/gau116

Ziegler, B., Schlosser, M., Lühder, F., Strebelow, M., Augstein, P., Northemann, W., Powers, A., & Ziegler, M. (1996). Murine monoclonal glutamic acid decarboxylase (GAD)65 antibodies recognize autoimmune-associated GAD epitope regions targeted in patients with type 1 diabetes mellitus and Stiff-man syndrome. Acta Diabetologica, 33(3), 225–231. https://doi.org/10.1007/bf02048548

